# VPS4 and CHMP7 release centromeres from the nuclear envelope for post-mitotic positioning in daughter nuclei

**DOI:** 10.64898/2026.07.13.738224

**Authors:** Nikolay Kornakov, Tyler Heiss, Arielle Kolodzinski, Rebecca Tam, Megan E. Kelley, Natalie Jones, Amalia H. Pasolli, Tarun M. Kapoor

## Abstract

Eukaryotic chromosomes occupy ordered configurations within the nucleus, an organization that must be re-established in daughter cells as the nuclear envelope reforms at the end of mitosis. The conserved enzyme VPS4 and ESCRT-III proteins mediate nuclear envelope reformation, yet their role in post-mitotic centromere positioning remains unclear. Here, we develop a chemical genetics approach to analyze the role of VPS4 in human cells. VPS4 inhibition prevents the clearance of CHMP7 from centromeres, which remain constrained in ring-like configurations established during mitosis. Without VPS4 activity, CHMP7, but not other ESCRT-III proteins, forms nuclear foci, nuclear envelope protein distribution is altered and inner nuclear membrane invaginations appear. Following these defects, DNA damage is observed in the vicinity of centromeres. Depletion of CHMP7, but not CHMP4B, suppresses this damage. We propose that VPS4-mediated turnover of CHMP7 releases centromeres from transient nuclear envelope contacts, ensuring their proper positioning after mitosis and maintaining genome integrity.

## INTRODUCTION

Eukaryotic chromosomes adopt spatial order within the nucleus, with specific chromosomal regions occupying defined positions relative to one another and to the nuclear envelope^1,2,3^. Recent studies have shown that the positioning of many genomic loci depends on DNA replication, and the distribution of centromeres in interphase nuclei relies on orderly progression through mitosis^4,5^. Establishing this organization in daughter cells requires a coordinated remodeling of chromatin-nuclear envelope interactions during nuclear envelope reassembly at the end of mitosis^6–8^. Centromeres present a distinct structural and temporal challenge to this process, as they remain attached to microtubules through late anaphase, creating a physical barrier that delays local membrane closure^9,10^. The mechanisms that mediate nuclear envelope sealing at these sites have been characterized^9–11^, however, how centromere positioning is coordinated with nuclear envelope reformation remains unknown.

Nuclear envelope gaps at mitotic exit are sealed by the Endosomal Sorting Complex Required for Transport-III (ESCRT-III) and its associated AAA (ATPases associated with diverse cellular activities) mechanoenzyme VPS4^12–15^. Human cells have twelve genes encoding ESCRT-III subunits (CHarged Multivesicular body/CHromatin-Modifying Proteins -CHMPs) and two VPS4 paralogs, VPS4A and VPS4B^13,16^. CHMPs polymerize at membrane gaps, and promote membrane sealing in a VPS4-dependent manner^11,17,18^. In current models, nuclear envelope integrity defects are detected by CHMP7 and LEMD2, its binding partner at the inner nuclear membrane^9,10,19,20^. Sealing gaps in the nuclear envelope establishes nuclear-cytoplasmic compartmentalization after chromosome segregation and is accompanied by the separation of LEMD2 from CHMP7, achieved in part by the export of CHMP7 from the nucleus^21,22,23^. LEMD2 remains nuclear, and together with other LEM-domain proteins and their binding partners BAF and Lamin A/C, mediates anchoring of the nuclear envelope to chromatin^8^, a set of interactions that must be remodeled for chromosomes to reach their interphase nuclear positions. However, whether VPS4/ESCRT-III membrane-sealing activity at mitotic exit is coupled to the remodeling of these chromatin-nuclear envelope contacts to establish chromosome organization in daughter cells is not known.

Support for the possibility of such coupling in human cells comes from studies of the fission yeast *Schizosaccharomyces japonicus*, which undergoes partial nuclear envelope breakdown during mitosis^24,25^. In this organism, CMP7 (the CHMP7 ortholog) in complex with Lem2 (the LEMD2 ortholog) recruits VPS4/ESCRT-III to disassemble the chromatin-bound Lem2-Nur1 complex, releasing heterochromatin from the inner nuclear membrane in a step required for chromosomes release from the nuclear envelope at mitotic entry^26^. Nur1 has no known metazoan ortholog. We note that in human cells, LBR (Lamin B Receptor), an inner nuclear membrane protein that binds both lamins and heterochromatin, and LEM-domain proteins tether heterochromatin to the nuclear periphery, and their combined loss causes global repositioning of heterochromatin, and notably centromeres toward the nuclear interior^27,28^. How these tethers release centromeres from the nuclear envelope at the end of mitosis in human cells remains unknown.

Acute inhibition of VPS4 in human cells would allow analyses of its functions in dynamic cellular processes such as mitosis and centromere positioning. Because VPS4 enzyme activity is essential^29^, the use of approaches that require days to take an effect, such as inducible CRISPR/Cas9-mediated gene knockout or shRNA-mediated depletion, or overexpression of dominant-negative enzyme mutants, as used in previous studies are complicated by loss of cell viability and the accumulation of pleiotropic phenotypes^29,9–11,19,30^. Acute inhibition of VPS4 enzymatic activity using cell permeable chemical probes could provide a powerful alternative, yet no selective and potent chemical inhibitors of VPS4 have been reported, and the available compounds lack specificity, inhibiting other AAA ATPases (e.g. the unfoldase VCP/p97)^31–33^. Chemical genetics has proven to be a powerful approach for dissecting functions of kinases and many other enzymes^34–36^, and more recently has been extended to AAA enzymes^37^. However, a chemical genetics system to analyze VPS4 functions in human cells has not been developed.

Recent studies identified paralog-specific VPS4 dependency as a therapeutically actionable vulnerability in more than 30% of human cancers^38,29,39^. This paralog-based synthetic lethality arises from the single-copy loss of *VPS4A* or *VPS4B* genes, which are positioned proximal to the tumor suppressors genes *CDH1* (on chromosome *16q)* or *SMAD4 (*on chromosome *18q*), respectively, and are frequently co-deleted. In these cancers, the remaining VPS4 paralog becomes essential for cell survival, as demonstrated by shRNA mediated depletion and CRISPR/Cas9 knockout experiments^29,39^. However, it is unclear whether paralog-selective inhibition of VPS4 enzyme activity would have effects similar to those due to protein depletion and whether such inhibition would selectively target cancer while sparing normal cells.

Here, we develop and employ a chemical genetics approach that allows acute and selective inhibition of VPS4 in human cells. Using this system, we uncover a role for VPS4 and its substrate CHMP7 in remodeling nuclear envelope-centromere contacts established during nuclear envelope reassembly at mitotic exit. We find that VPS4 activity is required for centromeres to reach their proper positions within the interphase nucleus. In the absence of VPS4 activity, CHMP7 is not cleared from chromatin, persisting at a subset of centromeres as well as other nuclear sites, with the centromeres remaining in the ring-like configurations established by the mitotic spindle. This failure to release centromeres is followed by DNA damage in their vicinity, which is suppressed upon depletion of CHMP7, but not CHMP4B or other ESCRT-III proteins. Our findings suggest that VPS4 and CHMP7 are part of a complex required to remodel and release LEM-domain proteins from chromatin, detaching chromatin from the nuclear envelope and allowing its proper positioning within each daughter nucleus.

## RESULTS

### Characterizing the localization dynamics and essentiality of VPS4 paralogs and ESCRT-III proteins

To analyze the function of VPS4/ESCRT-III proteins during cell division in HeLa Flp-In T-Rex cells (hereafter, HeLa) we examined their localization. We generated HeLa cells expressing individual fluorescently tagged constructs of VPS4A, CHMP7 and CHMP4B. Consistent with earlier observations^19,11,9^, VPS4 and these ESCRT-III proteins transiently localized to segregating chromosomes during anaphase (∼10 minutes after anaphase onset, duration: ∼5 min; Figure 1A). The inner nuclear membrane protein LEMD2 was recruited to anaphase chromosomes with timing similar to that of CHMP7 and CHMP4B, but persisted through the remaining time of image acquisition. Recruitment of ESCRT-III subunits was accompanied by restoration of nuclear compartmentalization, as revealed by the nuclear accumulation of the reporter 3xmNG-NLS (nuclear localization signal; Figure 1A).

**Figure 1.**
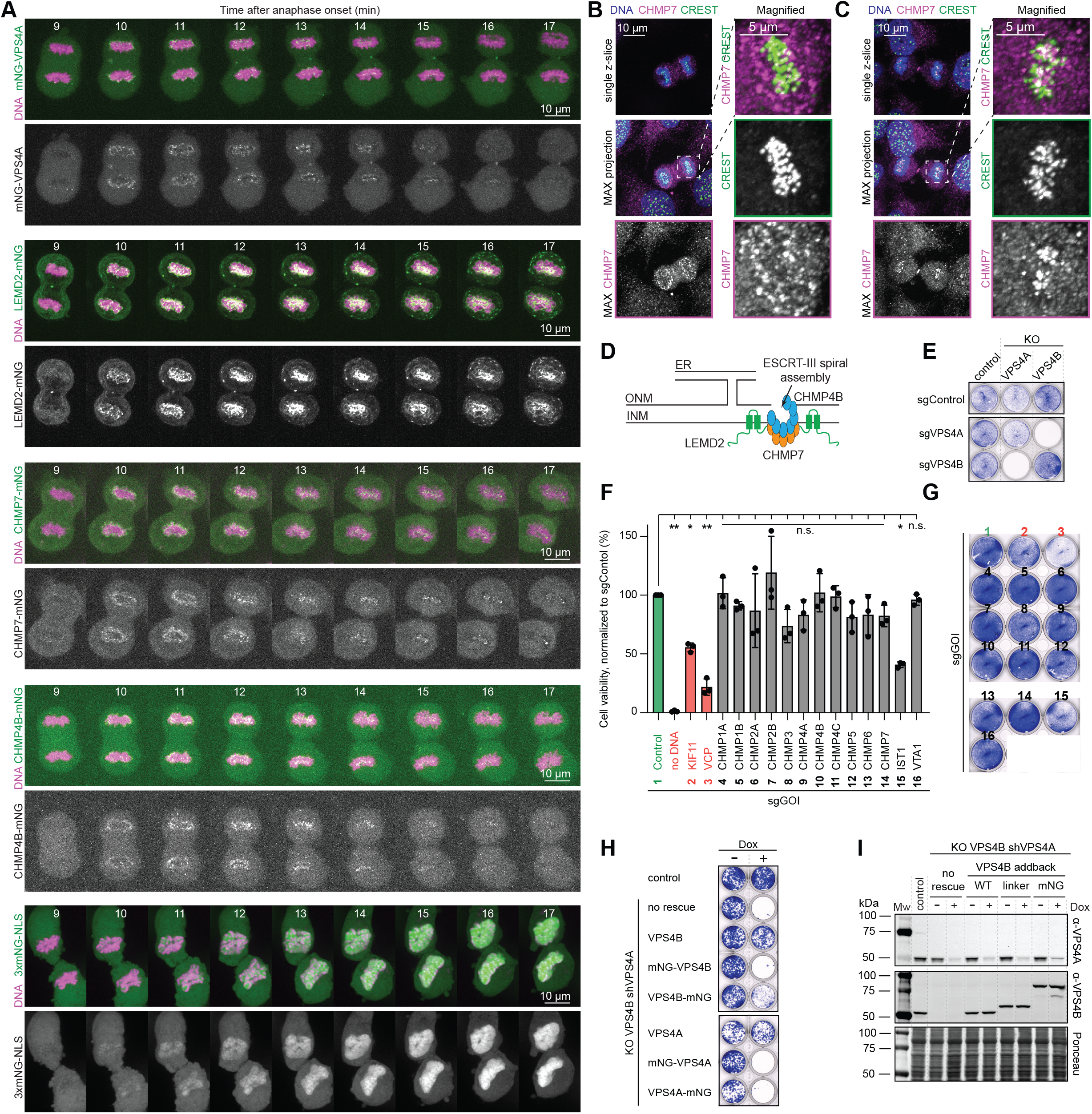
Characterizing the localization dynamics and essentiality of VPS4 paralogs and ESCRT-III proteins. (A) Representative time-lapse microscopy images of HeLa cell lines progressing through mitosis. Movies are temporally aligned between cells to the start of anaphase (t = 0). Cells express mNG-tagged constructs; ∼20 cells were imaged per condition. Maximum intensity projections of z-stacks are shown. Scale bars, 10 µm. (B-C) Immunofluorescence images of CHMP7 and centromeres (CREST) during nuclear envelope reformation. Single z-slice and maximum intensity projection are shown. Magnified boxed region is also shown. From top to bottom: merged image, CREST channel, CHMP7 channel. Scale bars are 10 µm and 5 µm, as indicated. (D) Schematic of VPS4/ESCRT-III recruitment to nuclear envelope gaps by LEMD2-CHMP7. Abbreviations: ER, endoplasmic reticulum; INM, inner nuclear membrane; ONM, outer nuclear membrane. (E) Analysis of cell viability following CRISPR-mediated depletion of one VPS4 paralog in cells carrying a monoclonal knockout of the other. Cells were fixed and stained (Crystal violet; 48 h post-selection of transfected cells). Control indicates parental HeLa cells; sgControl is an sgRNA targeting a ‘gene desert’ on chromosome 2. (F) Analysis of cell viability following CRISPR-mediated depletion of ESCRT-III subunits and VTA1 (a VPS4 cofactor) using 96-well plate assays. Cell viability was quantified and normalized to that of sgControl-transfected cells (100%). (Resazurin assay; 48 h post-selection of transfected cells). As controls, KIF11 (kinesin-5) and VCP/p97, two essential genes required for mitosis and proteastasis, respectively, were also depleted (red). Numbers under gene names refer to panel G. Abbreviation: GOI, gene-of-interest. Mean values ± SD are shown (n = 3, 3 technical repeats each). One way ANOVA with Dunnett’s multiple comparisons test compared to sgControl (green). **p<0.0001; *p<0.01; n.s., not significant. (G) Analysis of cell viability following depletion of individual ESCRT-III subunits. Cells were fixed and stained (Crystal violet; 48 h post-selection of transfected cells). Numbers refer to sgRNAs targeting genes, as in panel F. (H) Analysis of cell viability in cells expressing mNG-tagged VPS4A and VPS4B constructs as the only source of VPS4 in cells. Depletion of endogenous VPS4A was induced by addition of doxycycline (Dox, 2 µg/mL; cells were grown for 8 days). Corresponding immunoblot analysis is in (I). (I) Immunoblot analysis of KO VPS4B shVPS4A cells rescued with VPS4B constructs expressed at near-endogenous levels (compared to Control). Depletion was induced by addition of Dox (2 µg/mL for 48 h). See also Figure S1.

We noted that during the recruitment phase, VPS4/ESCRT-III foci first appeared on the spindle-distal (non-core) surface of the segregating chromosome masses, then dispersed, and subsequently reappeared on the spindle-facing (core) surface^40–42^, in the region occupied by centromeres and their attached microtubules (Figure 1A). Previous studies have reported localization of the LEMD2-CHMP7 complex at kinetochore microtubules in the vicinity of centromeres during nuclear envelope reformation^9,10^. To confirm that these inner foci correspond to centromeres, we performed immunofluorescence, co-staining against CHMP7 and centromeres using CREST serum (an autoimmune serum from patients with systemic sclerosis, that recognizes centromere proteins). CHMP7 staining on chromosomes was observed in a fraction of cells in anaphase. Earlier in anaphase, when the segregating chromosomes were still close together, CHMP7 was largely restricted to the nuclear rim. Later in anaphase, when the chromosomes had moved farther apart, CHMP7 was found at centromeres and was no longer detected at the rim (Figure 1B and 1C).

Given the involvement of VPS4/ESCRT-III in nuclear envelope reformation (Figure 1A and 1D) and transient recruitment to centromeres (Figure 1C) during this process, we examined which of these proteins were essential in HeLa cells, before initiating functional analyses. Consistent with earlier studies, knockout of individual VPS4 paralogs resulted in viable HeLa cells (hereafter, KO VPS4A and KO VPS4B), while knockout of both VPS4 paralogs was lethal^29,39^ (Figure 1E and S1A). In addition, we find that most individual ESCRT-III subunits are dispensable for HeLa cell survival (Figure 1F, 1G and S1B). The depletion of IST1 (CHMP8), one of the regulators of VPS4^43–45^, reduced HeLa cell viability (Figure 1F and 1G). We note, that DepMap datasets (https://depmap.org/portal)^46^ indicate that only three ESCRT-III subunits are essential across a majority of the other cell lines examined (Chronos score: ≤ −1 for CHMP2A, CHMP4B, and CHMP6; Figure S1C). As a subset of ESCRT-III subunits have paralogs, we co-depleted paralog pairs. CHMP1A/B or CHMP2A/B co-depletion reduced growth but cells remained viable (Figure S1D-S1G). These results indicate a complex pattern of functional redundancy within the ESCRT-III family. The dispensability of CHMP7, the only known recruiter of ESCRT-III proteins to the nuclear envelope^9,19,47^, indicates functional redundancy with other nuclear envelope sealing pathways, and contrasts with the essential role of VPS4 activity, which is retained as long as one paralog is present.

Therefore, we first chose to target VPS4 to study consequences of its inactivation in HeLa cells. As cells lacking both paralogs are not viable, we established a conditional depletion system in which the knockdown of one VPS4 paralog could be examined in cell lines with the other paralog knocked out (Figure S1H-S1O). We found that ∼50% reduction in VPS4A levels in KO VPS4B cell lines resulted in loss of cell viability^29^ and the accumulation of DNA damage (Figure S1L-S1N), consistent with previous findings^9^. Further, achieving ∼90% knockdown required ∼48 hours, with the protein depletion beginning at ∼24 hours after induction, a time course that limits interpretations of phenotypes in dynamic processes such as cell division (Figure S1O).

An alternative approach to study VPS4 function would be the use of an inducible degron system, such as an AID tag^48^. However, attempts to introduce AID or fluorescent tags on VPS4A and VPS4B using CRISPR/Cas9, making these tagged constructs the sole source of VPS4 in the cells, did not yield viable clones. We hypothesized that tagging VPS4 could compromise its function. Therefore, we used our conditional depletion system to deplete endogenous VPS4 and replace it with untagged, fluorescently tagged, or degron-tagged VPS4 alleles expressed at near-endogenous levels (Figure 1H, 1I, and S1P-S1R). Untagged VPS4A and VPS4B were able to rescue lethality upon depletion of endogenous VPS4A, while N- and C-terminally mNeonGreen (mNG) tagged constructs failed to do so. Similarly, N-terminal AID tagging of VPS4A did not yield viable cells. Of the tagged construct tested, the C-terminally tagged VPS4B with an extended Gly-Ser linker between VPS4B and mNG was the least compromised. However, it remained substantially impaired (Figure 1H), supporting our conclusion that tagging VPS4 disrupts its function and motivating the development of a tag-free strategy.

Together, these data indicate that analyzing the cell division functions of VPS4 and ESCRT-III, proteins with dynamic localizations and functional redundancies, will require a different approach.

### A chemical genetics system to examine VPS4 function

We next focused on employing a chemical genetics approach to analyze VPS4 function in HeLa cells. Recent studies have shown that an engineered cysteine mutation in VPS4 can confer sensitivity to ASPIRe-1, a designed covalent chemical inhibitor that does not inhibit wildtype VPS4, or other related AAA proteins, in biochemical assays^37^. In particular, VPS4B D135C was found to be sensitive to the compound, but this mutation altered enzymatic parameters ∼6-fold^37^. We introduced VPS4 with ASPIRe-1-sensitising cysteine mutations into our conditional depletion system but found that these mutant alleles did not fully support cell growth (Figure 2A and 2B), likely due VPS4’s reduced ATPase activity.

**Figure 2.**
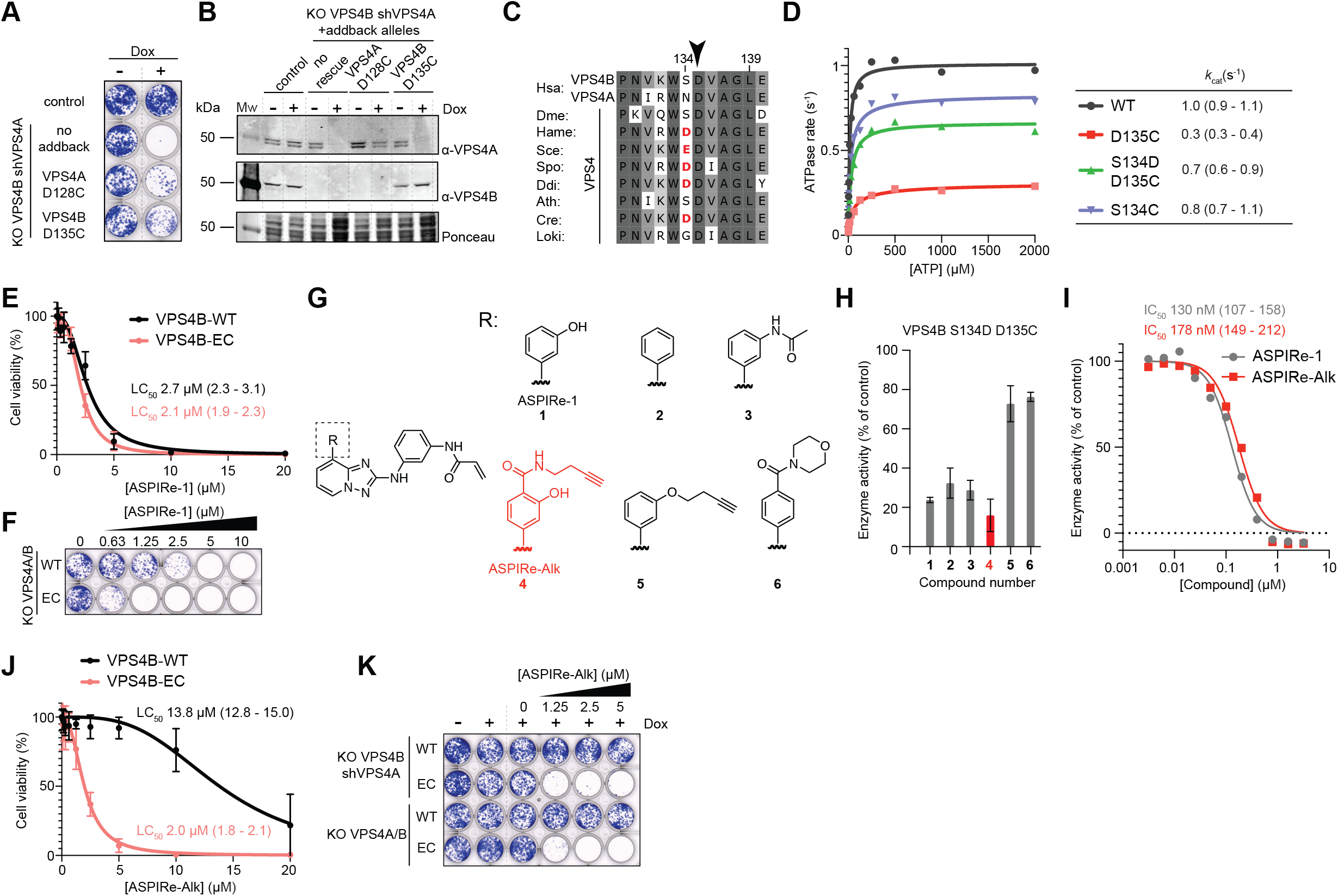
Developing a chemical genetics system to examine VPS4 function. (A) Analysis of ASPIRe-1-sensitive alleles of VPS4A and VPS4B in a VPS4 conditional depletion system. Depletion was induced by addition of Dox (2 µg/mL; cells were grown for 8 days). Control indicates parental HeLa cells. (B) Immunoblot analysis of KO VPS4B shVPS4A cells rescued with shRNA-resistant VPS4A-D128C or VPS4B-D135C. Depletion was induced by addition of Dox (2 µg/mL for 48 h). (C) Sequence alignment of VPS4 orthologs. The position of the cysteine mutation is indicated (Asp135 in human VPS4B, black arrow). A negatively charged amino acids preceding Asp135 residue are highlighted (red). Organism codes follow KEGG nomenclature (https://www.genome.jp/kegg/). (D) Analysis of enzymatic parameters of VPS4B-WT and mutants using an NADH-coupled ATPase assay. Dots represent averages of two experiments, and lines are fits for allosteric sigmoidal enzyme kinetics. Values for k_cat_ were estimated from the fits and are shown at the right (95% CI are shown in parenthesis). Hill coefficients (*h*) and R^2^: for VSP4B-WT (1.03; R^2^=0.90), for VPS4B-D135C (0.73; R^2^=0.95), for VPS4B-DC (1.01; R^2^=0.83), for VPS4B-S134C (0.83; R^2^=0.87). (E) Analysis of cell viability following ASPIRe-1 treatment (72 h; 96-well plate assays). VPS4B-WT (black) and VPS4B-EC (red) are shown. Values were normalized to DMSO-treated conditions (100%). Points represent mean values ± SD (n = 3, 3 technical repeats each). LC_50_ with 95% CI are shown in the parentheses. Values are estimated based on a sigmoid inhibition curve fit with variable slope. Hill coefficients (*h*) and R^2^: for VSP4B-WT (−2.44; R^2^=0.96), for VSP4B-EC (−2.85; R^2^=0.98). (F) Colony formation assay of VPS4B-WT and VPS4B-EC cells treated by ASPIR-1; cells were grown for 8 days. (G) Chemical structures of ASPIRe-1 (compound 1) and a few analogs, ASPIRe-Alk (compound 4) is highlighted in red. (H) VPS4B-DC ATPase activity upon treatment by ASPIRe-1 analogs (3 µM, 30 min, 1 mM of ATP). ASPIRe-Alk is highlighted in red. Residual ATPase activity is shown. Bars are averages of four experiments with 95% SD. (I) VPS4B-DC treatment with ASPIRe-1 (gray) or ASPIRe-Alk (red); 30 min pre-treatment time (at 1 mM of ATP). IC_50_ values with 95% CI are shown in parenthesis. IC_50_ were estimated based on sigmoid inhibition curve fits. Hill coefficients (*h*) and R^2^: for ASPIRe-1 (−1.89; R^2^=0.99), for ASPIRe-Alk (−1.83; R^2^=0.97). (J) Analysis of cell viability of VPS4B-WT (black) and VPS4B-EC (red), (72 h after ASPIRe-Alk addition; 96-well plate assays). Values were normalized to DMSO-treated conditions (100%). Points represent mean values ± SD (n = 3, 3 technical repeats each). LC_50_ with 95% CI are shown in parentheses. Values were estimated based on sigmoid inhibition curve fits with variable slope. Hill coefficients (*h*) and R^2^: for VSP4B-WT (−3.27; R^2^=0.94), for VSP4B-EC (−2.44; R^2^=0.99). (K) Colony formation assay examining allele-specific viability upon treatment of VPS4B-EC or VPS4B-WT expressing cells by ASPIRe-Alk in the VPS4 conditional depletion background and in CRISPR-engineered cells (8 days). See also Figure S2.

To use this chemical genetics approach we focused on VPS4B and examined secondary mutations that could restore ATPase activity and cellular function. We noted that in some species that express a single VPS4 gene a negatively charged amino acid precedes the conserved Asp-135 residue. In addition, we found that there is a charged residues at equivalent site in other AAA proteins in the meiotic clade (Figure 2C and S2A). We generated recombinant forms of VPS4B with an Asp residue adjacent to the sensitizing Cys residue (i.e. S134D in addition to D135C) and found it to be an active ATPase (Figure 2D and S2B). Gratifyingly, our conditional depletion system indicated that double mutants carrying the D135C mutation together with either the S134D or S134E mutation are viable and sensitive to ASPIRe-1 (Figure S2C and S2D). We also tested VPS4B S134C, which moved the sensitizing Cys residue to the 134 position and found this to a biochemically active enzyme that was functional in cells. However, consistent with the inhibitor binding pose, this allele was not sensitive to ASPIRe-1 (Figure 2D, S2C, and S2D).

Next, to generate a cell line expressing the ASPIRe-1 sensitive VPS4B allele as the only source of VPS4, we introduced the double mutants (S134D D135C or S134E D135C) into the cells with VPS4B knocked out, and then also knocked out VPS4A. These cells were viable (Figure S2E and S2F), and we were able to isolate S134E D135C VPS4B (hereafter VPS4B-EC) and VPS4B-WT control clones. Dose-dependent analyses revealed only a ∼1.3-2-fold difference in the sensitivity of these clones to ASPIRe-1 (LC_50_ values for VPS4B-WT: 2.7 µM, VPS4B-EC: 2.1 µM in short-term assays; 0.6 µM of ASPIRe-1 suppressed colony formation of VPS4B-EC cells; Figure 2E and 2F). We hypothesized that this limited difference in potency between VPS4B-WT and VSP4B-EC cell lines was due to the compound’s off-target activity.

We hypothesized that the inhibition of kinases, which are known targets of compounds with the triazolopyridine scaffold, are likely the source of VPS4-inhibition independent toxicity^37,49,50^. To address that, we used our structural data for the ASPIRe-analogs bound to VPS4B complex (PDB: 7L9X) and available kinase binding data for the scaffold, to design a handful of compounds. We focused on triazolopyridine analogs with 5’-aryl substituents, as extending along this vector may retain VPS4B affinity while disrupting off-target interactions (Figure 2G). Consistent with this reasoning, we found ASPIRe analogs to be active against VPS4B-DC allele in biochemical assays (Figure 2H). One analog (hereafter, ASPIRe-Alk or Alk) is a potent inhibitor of VPS4B-DC that blocks activity in a time-dependent manner, consistent with covalent binding (IC_50_ ∼180 nM, incubation time: 30 min, Figure 2I and S2G).

We tested ASPIRe-Alk, along with two analogs in cells, and gratifyingly, VPS4B-EC was selectively sensitive to ASPIRe-Alk in both the VPS4 conditional depletion background and in CRISPR-engineered cells (∼7-fold difference between the LC_50_ values for VPS4B-WT: 13.8 µM and VPS4B-EC: 2 µM in short-term assays; 1.25 µM of Alk suppressed colony formation of VPS4B-EC cells; Figure 2J, 2K, S2H-S2J).

Together, these data show how structure-guided inhibitor modifications and secondary mutations can establish a chemical genetics system to acutely inhibit VSP4B in human cells.

### The chemical genetics approach allows acute inhibition of VPS4 activity in cells

To use ASPIRe-Alk to examine VPS4 function in cells, we first analyzed target engagement using the compound’s alkyne handle and click chemistry^51^. For these studies, we analyzed cells overexpressing either VPS4B-WT or cysteine mutant alleles (VPS4B-D135C, VPS4B-DC or VPS4B-EC). Cells were treated with increasing concentrations of the Alk compound (up to 5 µM, 4 h), followed by lysis and fluorophore conjugation. We observed a prominent labeled band in cells expressing VPS4B D135C and S134D D135C mutants, but not in cells expressing the wild-type protein or in non-transfected cells (Figure 3A). Labeling of VPS4B-EC by Alk (5 µM) was time-dependent and could be observed within 2 hours (Figure 3B). Co-treatment of cells with Alk and increasing concentrations of ASPIRe-1 suppressed specific labeling of the VPS4B cysteine allele, consistent with stoichiometric target occupancy, while a weak background band remained largely unchanged (Figure 3C).

**Figure 3.**
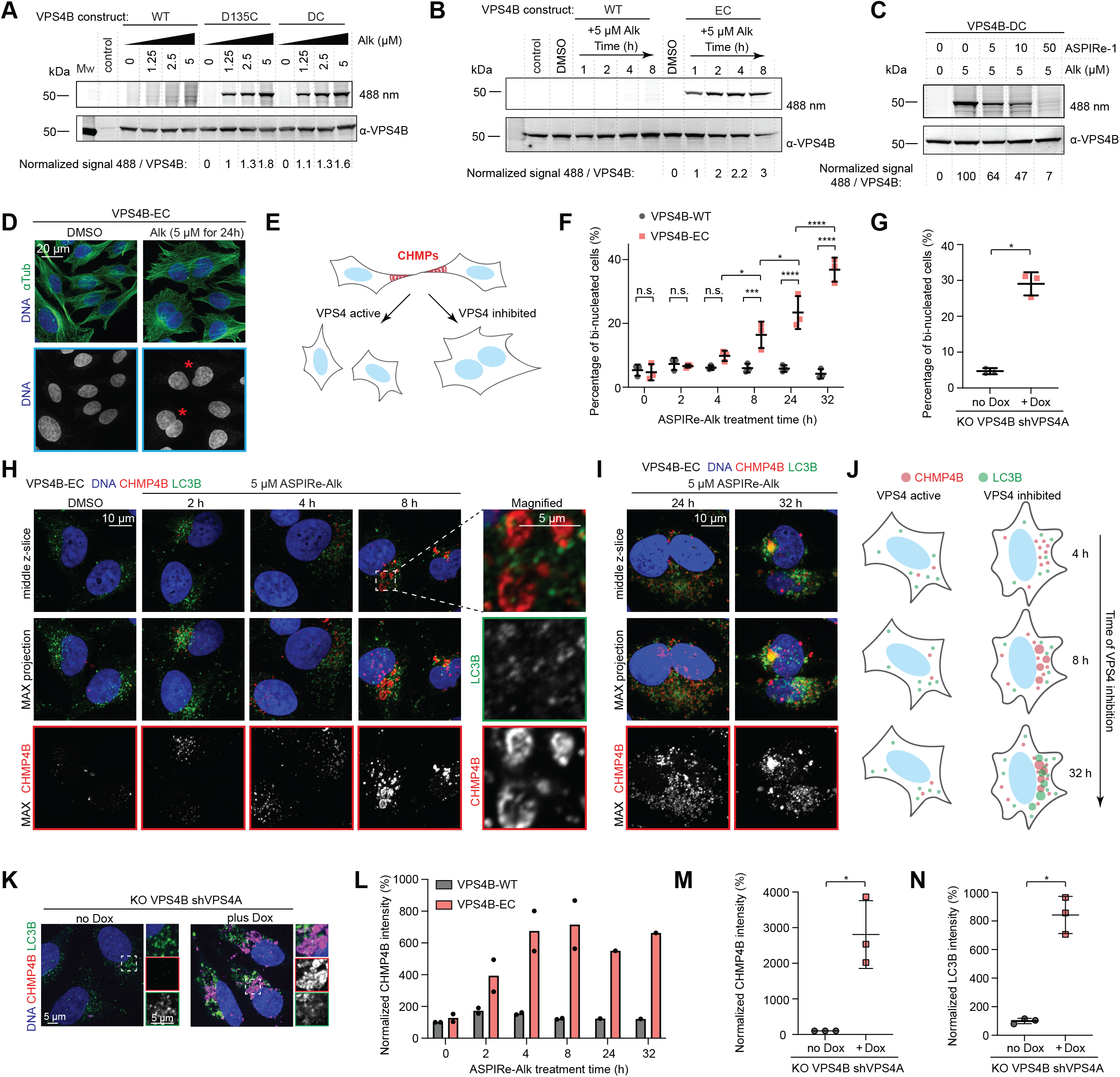
Chemical genetics approach allows acute inhibition of VPS4 activity in cells. (A) Analysis of engagement of VPS4B cysteine mutants by ASPIRe-Alk in cells. HEK293T cells, expressing the indicated VPS4B constructs were treated with different concentrations of ASPIRe-Alk (4 h), followed by lysis and fluorophore conjugation (click-chemistry). Control indicates non-transfected cells. Corresponding SDS-PAGE in-gel fluorescence (488 nm) and immunoblot are shown. The normalized 488 nm / VPS4B signal ratio is shown for VPS4B cysteine mutants; the ratio at the 1.25 µM of ASPIRe-Alk was set to 1. (B) Analysis of time-dependence of VPS4B-EC engagement by ASPIRe-Alk in cells. HEK293T cells expressing the indicated VPS4B constructs were treated with ASPIRe-Alk, followed by lysis and fluorophore conjugation (click-chemistry). Control indicates untreated cells. SDS-PAGE in-gel fluorescence (488 nm) and immunoblot analysis are shown. The normalized 488 nm / VPS4B signal ratio is shown for VPS4B-EC. The ratio for 1 h of ASPIRe-Alk treatment was set to 1. (C) Analysis of competition between ASPIRe-1 and ASPIRe-Alk for VPS4B cysteine mutant engagement. HEK293T cells expressing VPS4B-DC were co-treated with both compounds (indicated concentrations for 4 h 30 min), followed by lysis and fluorophore conjugation (click-chemistry). SDS-PAGE in-gel fluorescence (488 nm) and immunoblot analysis are shown. The normalized 488 nm / VPS4B signal ratio is shown. The ratio for 0 µM of ASPIRe-1 treatment was set to 100. (D) Representative immunofluorescence images of cells upon VPS4B-EC treatment with ASPIRe-Alk (5 µM for 24 h). Bi-nucleated cells are indicated (asterisk). Alpha-Tubulin and DNA staining were used to identify bi-nucleated cells. Maximum intensity projections of z-stacks are shown. Scale bar, 20 µm. (E) Schematic of the role of VPS4/ESCRT-III in cell abscission. VPS4 inactivation leads to cytokinetic failure and formation of bi-nucleated cells. (F) Time-dependent accumulation of bi-nucleated cells upon VPS4B-EC cells treatment with ASPIRe-Alk; the fraction of bi-nucleated cells is plotted. Mean values ± SD are shown (3 independent experiments; total number of cells per time point for VPS4B-WT: 434; 554; 629; 436; 499; 571; for VPS4B-EC: 473; 525; 640; 588; 470; 113). One way ANOVA with Šídák’s multiple comparisons test. ****p <0.0001; ***p= 0.0005; *p<0.05; n.s., not significant. (G) Analysis of the accumulation of bi-nucleated cells upon VPS4 depletion induced by Dox addition (2 µg/mL for 48 h); the fraction of bi-nucleated cells is plotted. Mean values ± SD are shown (3 independent experiments; total number of cells for no Dox: 586; for +Dox: 328). Unpaired t test; *p=0.0002. (H) Schematic of the role of VPS4/ESCRT-III in vesicle transport. VPS4 inhibition leads to perinuclear accumulation of CHMP4B, followed by LC3B accumulation with an ∼24 h delay relative to CHMP4B. (I) and (J) Immunofluorescence images of CHMP4B and LC3B upon VPS4B-EC cells treatment with ASPIRe-Alk (5 µM; indicated time). Middle z-slice and maximum intensity projection are shown; scale bar, 10 µm. Magnified boxed region is also shown; scale bar, 5 µm. (K) Immunofluorescence images of CHMP4B and LC3B accumulation upon VPS4 depletion, induced with Dox (2 µg/mL for 48 h). Magnified boxed region is also shown. From top to bottom: merged image, CHMP4B channel, LC3B channel. Scale bars, 5 µm. (L) Analysis of CHMP4B accumulation following VPS4B-EC cells treatment with ASPIRe-Alk (5 µM; indicated time). Average values of two experiments, with two technical repeats each are shown as bars; dots are mean values of each of two datasets. (M) Analysis of CHMP4B accumulation following VPS4 depletion induced with Dox (2 µg/mL for 48 h). Values were normalized to the no Dox condition (100%). Mean values ± SD are shown (3 independent experiments; total number of cells for no Dox: 448; for +Dox: 245). Unpaired t test; *p=0.0079. (N) Analysis of LC3B accumulation following VPS4 depletion induced with Dox (2 µg/mL for 48 h). Values were normalized to the no Dox condition (100%). Quantification was performed using part of the dataset, used in panel M. (total number of cells for no Dox: 319; for +Dox: 166). Mean values ± SD are shown. Unpaired t test; *p=0.0006. See also Figure S3.

To compare phenotypes resulting from acute inhibition of VPS4 with those caused by VPS4 protein depletion, we first focused on cytokinesis^52^ and endosomal transport^53^, well-characterized VPS4/ESCRT-III functions. Cells were treated with ASPIRe-Alk and analyzed by immunofluorescence (Figure 3D). In VPS4B-EC cells, Alk treatment led to a time-dependent accumulation of bi-nucleated cells, consistent with failure in cytokinetic abscission (Figure 3D, 3E, and 3F). The effect was evident at 8 hours (∼16% bi-nucleated cells) and strengthened by 32 hours (∼37% of cells), at which point cell number decreased and cell death was observed. Importantly, Alk-treated VPS4B-WT cells did not show an increase in the fraction of bi-nucleated cells (Figure 3F). In addition, we also observed a similar phenotype using our VPS4 conditional depletion system (bi-nucleated cells: no Dox **∼**5%, plus Dox **∼**29%; Figure 3G).

We next examined the time-dependent accumulation of CHMP4B, a major ESCRT-III subunit^9,54^, and LC3B, an autophagy marker^55^, upon VPS4B inhibition (Figure 3H). In VPS4B-EC cells, CHMP4B foci began to appear after 2 hours of Alk addition, increasing in intensity and size over ∼8 hours (Figure 3H-3J and 3L). In addition to forming foci, CHMP4B localized to vesicle-like objects (Figure 3H), with most of the signal accumulating in the perinuclear region overlapping with the Golgi compartment (Figure S3C). In contrast, LC3B accumulation was not an immediate consequence of VPS4 inactivation. Increase in LC3B puncta intensity and area was detected only at the final time point (32 hours), revealing a **∼**24-hour lag relative to CHMP4B accumulation (Figure 3H-3J and S3D). Again, we did not observe these phenotypes in ASPIRe-Alk-treated VPS4B-WT control cells (Figure S3A) and recapitulated them in our VPS4 conditional depletion system (Figure 3K, 3M, and 3N). We note that at the 8 h time point, there was little overlap between CHMP4B and LC3B signals (Figure 3I, 3K, and S3B), with LC3B occupying more peripheral regions of the cytoplasm than CHMP4B.

Together, these data indicate that our chemical genetics system allows acute and selective inactivation of VPS4 and can be used to examine time-dependent phenotypes following loss of enzyme activity in human cells.

### VPS4 inhibition traps CHMP7 and LEM-domain proteins at centromeres after mitosis and disrupts nuclear envelope organization

Upon nuclear envelope reformation at the end of anaphase, the exposed chromatin surface forms numerous contacts with the reforming nuclear envelope^8,41^, at least a subset of which must be remodeled to restore nuclear architecture in daughter cells^6^. We find that CHMP7 is transiently recruited to chromatin and centromeres during this process and is subsequently cleared (Figure 1A-1C). Therefore, we analyzed the effects of VPS4 inhibition on the localization of CHMP7, LEMD2, the inner nuclear membrane binding partner of CHMP7, and Emerin, the related LEM-domain protein. To reduce phenotype variability, we examined synchronized cells. We arrested cells in S phase using a thymidine block and either released cells from the thymidine block and allowed them to progress through mitosis, or kept them arrested in G2 phase by inhibiting CDK1 with RO3306^56^. In both cases VPS4 was inhibited for a 18 h total (Figure 4A, 4B, and S4A-S4D).

**Figure 4.**
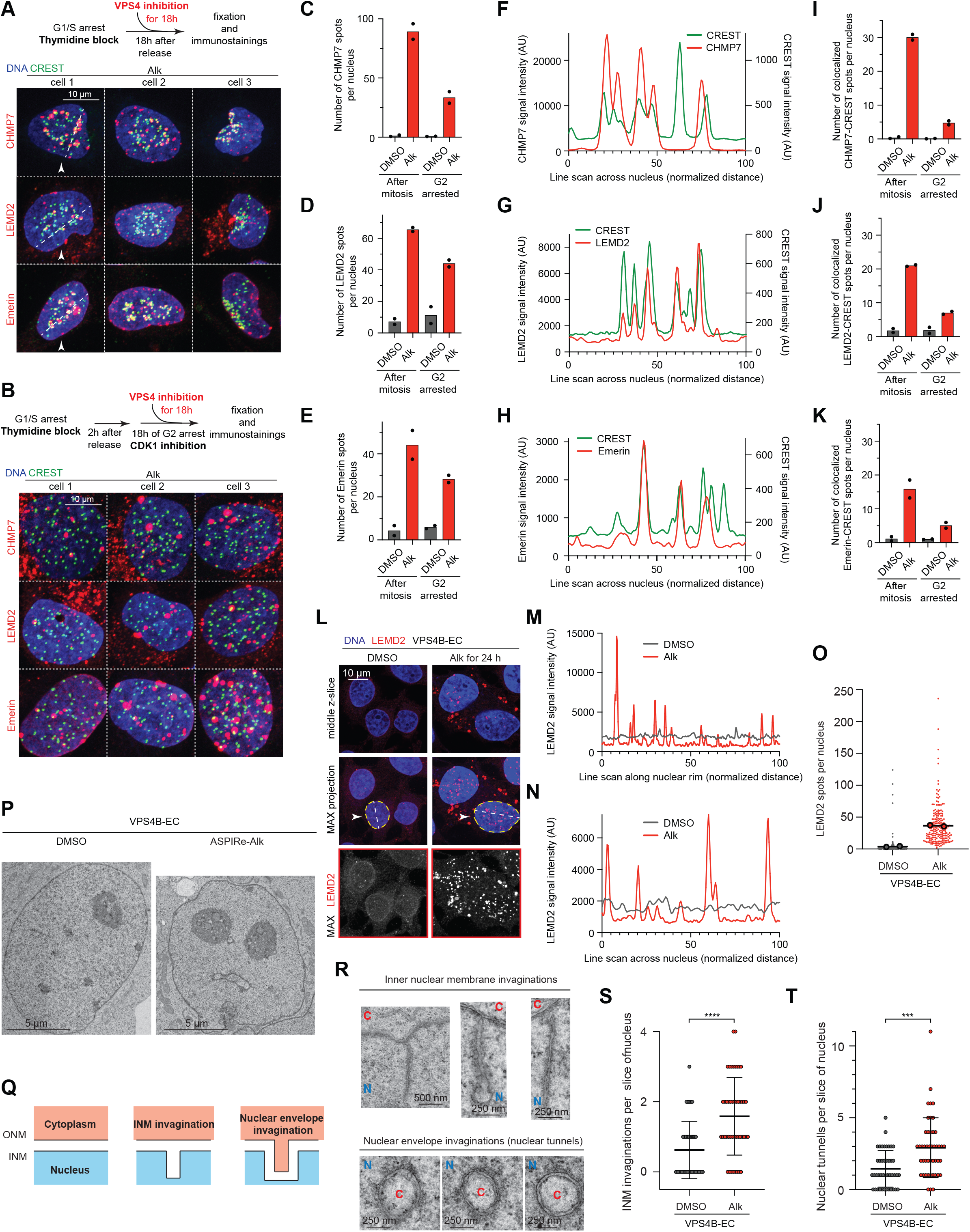
VPS4 inhibition traps CHMP7 and LEM-domain proteins at centromeres after mitosis and disrupts nuclear envelope organization. (A) and (B) Representative immunofluorescence images of CHMP7, LEMD2, Emerin, and centromeres following treatment of VPS4B-EC cells with ASPIRe-Alk (5 µM for 18 h). Cells were released from a thymidine block, and progressed through mitosis (A) or were arrested in G2 for 18h (B). Cells used for line scan analysis are indicated (arrowhead and dotted line). Maximum intensity projections of z-stacks are shown; scale bars, 10 µm. (C-E) Analysis of the number of CHMP7, LEMD2, or Emerin spots per nucleus following treatment of VPS4B-EC cells with ASPIRe-Alk (5 µM for 18 h) in post-mitotic and G2 arrested cells. Mean values of all cells are shown; dots are mean values of each of two datasets. (F-H) Line scan analysis (5 pxl width) of CHMP7, LEMD2, or Emerin (red) and CREST (green) signal intensity across the nucleus. (I-K) Analysis of the colocalization of CHMP7, LEMD2, or Emerin spots and CREST spots per nucleus following treatment of VPS4B-EC cells with ASPIRe-Alk (5 µM for 18 h) in post-mitotic and G2 arrested cells. Mean values are shown; the same dataset as in Figure 4C-4E was used for quantification. (L) Immunofluorescence images of LEMD2 in VPS4B-EC cells treated with DMSO or ASPIRe-Alk (5 µM for 24 h). Middle z-slices and maximum intensity projections are shown. Cells used for line scan analysis are indicated (arrowhead and dotted line). Scale bar, 10 µm. (M) and (N) Line scan analysis (5 pxl width) showing LEMD2 signal intensity along the nuclear rim and across the nucleus in DMSO (gray) and ASPIRe-Alk-treated cells (red). (O) Analysis of the number of LEMD2 spots per nucleus following treatment of VPS4B-EC cells with ASPIRe-Alk (5 µM for 24 h). Mean values of all data are shown; small dots represent individual nuclei; big dots are mean values of two datasets (total number of cells DMSO: 200; Alk: 211). (P) Serial section electron microscopy images of nuclei of VPS4B-EC cells treated with DMSO or ASPIRe-Alk (5 µM for 18 h). Scale bar, 5 µm. (Q) Schematic of the types of observed nuclear envelope alterations. (R) Magnifications of selected electron microscopy images revealing inner nuclear membrane (INM) invaginations and nuclear envelope invaginations. “N” indicates nucleus; “C” indicates cytoplasm. Scale bars are 500 nm and 250 nm. (S) and (T) Analysis of the number of INM invaginations and nuclear tunnels per slice of nucleus in VPS4B-EC cells treated with DMSO or ASPIRe-Alk (5 µM for 18 h). Mean values ± SD are shown (3 independent experiments; total number of nuclei for DMSO: 43 ; Alk: 46). Mann Whitney test; ****p<0.0001, ***p=0.0001 See also Figure S4.

Upon VPS4 inhibition, CHMP7, LEMD2, and Emerin formed foci in the nucleus, with more foci formed in cells after mitosis relative to the G2-arrested cells, with the biggest difference observed for CHMP7 (mean spots/nucleus for post-mitotic vs. G2-arrested Alk-treated cells: for CHMP7 ∼89 vs. ∼34, for LEMD2 ∼64 vs. ∼42, for Emerin: ∼38 vs. ∼27; Figure 4C-4E and S4A-S4D). In G2-arrested cells, VPS4 inhibition caused CHMP7, and to some extent LEMD2, to accumulate in the cytoplasm rather than the nucleus (Figure 4B). Because these cells have not undergone mitosis, this indicates that CHMP7 requires nuclear envelope breakdown to access chromatin.

We next examined the localization of CHMP7, LEMD2, and Emerin foci relative to centromeres in post-mitotic cells. CHMP7, LEMD2, and Emerin foci localized in the vicinity of centromeres (mean CREST-colocalized spots/nucleus in Alk-treated cells: for CHMP7 ∼30, for LEMD2 ∼21, for Emerin ∼13; Figure 4A and 4F-4K). Blocking cells from progression through mitosis diminished centromere colocalization for all three proteins (mean of CREST-colocalized spots/nucleus in Alk-treated cells: for CHMP7 ∼4, LEMD2 ∼7, Emerin ∼4; Figure 4B and 4I-4K). We found this post-mitotic centromere localization remarkable, although not every centromere was labelled with the examined proteins and not every CHMP7/LEMD2/Emerin foci was found at centromeres (Figure 4A).

In addition to post-mitotic centromere localization, we observed accumulation of LEMD2 and Emerin foci along the nuclear rim and within the nucleus in G2-arrested cells (Figure 4B), suggesting that VPS4 inhibition might disrupt the proteins anchoring chromatin to the nuclear envelope. LEMD2 and its binding partner Lamin A/C are proposed to be one of two mechanisms anchoring heterochromatin to the inner nuclear membrane (alongside LBR)^28,57^, however other LEM-domain proteins, including Emerin and LAP2, may act redundantly or cooperatively in this process^27^. Previous studies in fission yeast also suggested a possible role for VPS4/ESCRT-III in resolving attachments between the inner nuclear membrane and heterochromatin, dependent on the LEMD2 ortholog LEM2^26^. Thus, we further characterized the effect of VPS4 inhibition on LEM-domain proteins and Lamin A/C.

Under control conditions, LEMD2 displayed weak nuclear envelope localization and a diffuse cytoplasmic signal in interphase cells, with prominent accumulation in a fraction of micronuclei, as previously reported^23,58^ (Figure 4L and S4E). At late anaphase, we observed LEMD2 on chromatin, and following mitotic exit, it also formed cytoplasmic spots (Figure S4E and S4F). Upon VPS4 inhibition, LEMD2 formed numerous spot-like aggregates in the nucleus (mean spots/nucleus: DMSO ∼4, Alk ∼37; Figures 4L, 4N, and 4O), and coated vesicle-like objects in the cytoplasm. Some LEMD2 spots originated at the nuclear surface and extended to the nuclear interior, and LEMD2 showed heterogeneous, focally enriched distribution along the nuclear rim compared to control (Figure 4M). VPS4 inhibition produced similar phenotypes for Emerin and LAP2, as well as Lamin A/C. Emerin, LAP2, and Lamin A/C all formed intranuclear foci, with Emerin showing the most pronounced phenotype (mean of spots/nucleus for DMSO: ∼2, Alk-treated: ∼57; Figure S4G-S4P). The Emerin signal distribution along the nuclear rim also became more heterogeneous than control (Figure S4I), while the effects on LAP2 and Lamin A/C distribution were less pronounced (Figure S4L and S4O). Thus, loss of VPS4 activity disrupts the nuclear organization of LEM-domain proteins and Lamin A/C.

To determine whether changes in LEM-domain protein localization were accompanied by structural alterations of the nuclear envelope, we analyzed the effect of VPS4 inhibition on the nuclear envelope organization using serial section electron microscopy (Figure 4P). Guided by previous studies^59,60^, we classified nuclear membrane invaginations as either double-membrane structures (nuclear tunnels) or invaginations of the inner nuclear membrane alone. (Figure 4Q and 4R). In control cells, such structures were observed at low frequency, and upon VPS4 inhibition, the frequency of these invaginations increased, consistent with the fluorescence microscopy data (median number of tunnels/slice of nucleus for DMSO: 1, Alk: 3; for inner membrane invagination/slice of nucleus for DMSO: 0, Alk: 1.5; Figure 4S and 4T). In some cases, these invaginations originated from the inner membrane of nuclear tunnels and occasionally bridged adjacent tunnels. At the limit of our analysis, we did not detect obvious gaps in the nuclear envelope. We propose that foci of inner nuclear membrane proteins formed upon VPS4 inhibition correspond to nuclear tunnels and inner nuclear membrane invaginations observed by EM.

Together, these findings indicate that VPS4 clears CHMP7 and LEM-domain proteins from centromeres after mitosis and prevents the accumulation of inner nuclear membrane invaginations.

### VPS4 and CHMP7 ensure proper centromere positioning after mitosis

The persistence of CHMP7 and LEM-domain proteins at centromeres upon VPS4 inhibition raised the possibility that centromeres may not to be released from the nuclear envelope at the end of mitosis. During prometaphase, centromeres adopt a ring-like configuration, called the chromosome rosette^61,62^, with centromeres oriented towards spindle axis (Figure 5A). This chromosome organization is established by geometry of mitotic spindle^63^. During nuclear envelope reformation, centromeres cluster at the core region of the reassembling nuclear envelope and then redistributed within the nucleus as chromosomes decondense^64^ (Figure S5A and S5B). Consistent with our live-cell imaging of parental HeLa cells (Figure 1A-1C), CHMP7 in DMSO-treated VPS4B-EC cells coated chromatin during nuclear envelope reformation and cleared thereafter, with the most persistent signal at centromeres (Figure 1C and Figure S5B). Upon VPS4 inhibition, CHMP7 instead persisted on chromatin once recruited, with foci at the nuclear rim and at centromeres (Figure 5B and S5A). In control conditions, centromeres were distributed throughout the nucleus after chromosome decondensation, with some enrichment at nucleoli and depletion from the nuclear periphery^65,66^, and positioning was overall similar across post-mitotic, G2-arrested and asynchronous cells (Figure S4A and S4B). Notably, we observed altered centromere distribution following VPS4 inhibition after mitosis. The most common pattern was centromere clustering in ring-shaped configurations at the center of the nucleus; in a small subset of cells centromeres instead gathered at the nuclear periphery (Figure 5C and 4A). These patterns resembled the centromere clustering observed in prometaphase and anaphase in normal cells, the chromosome rosettes (Figure 5A).

**Figure 5.**
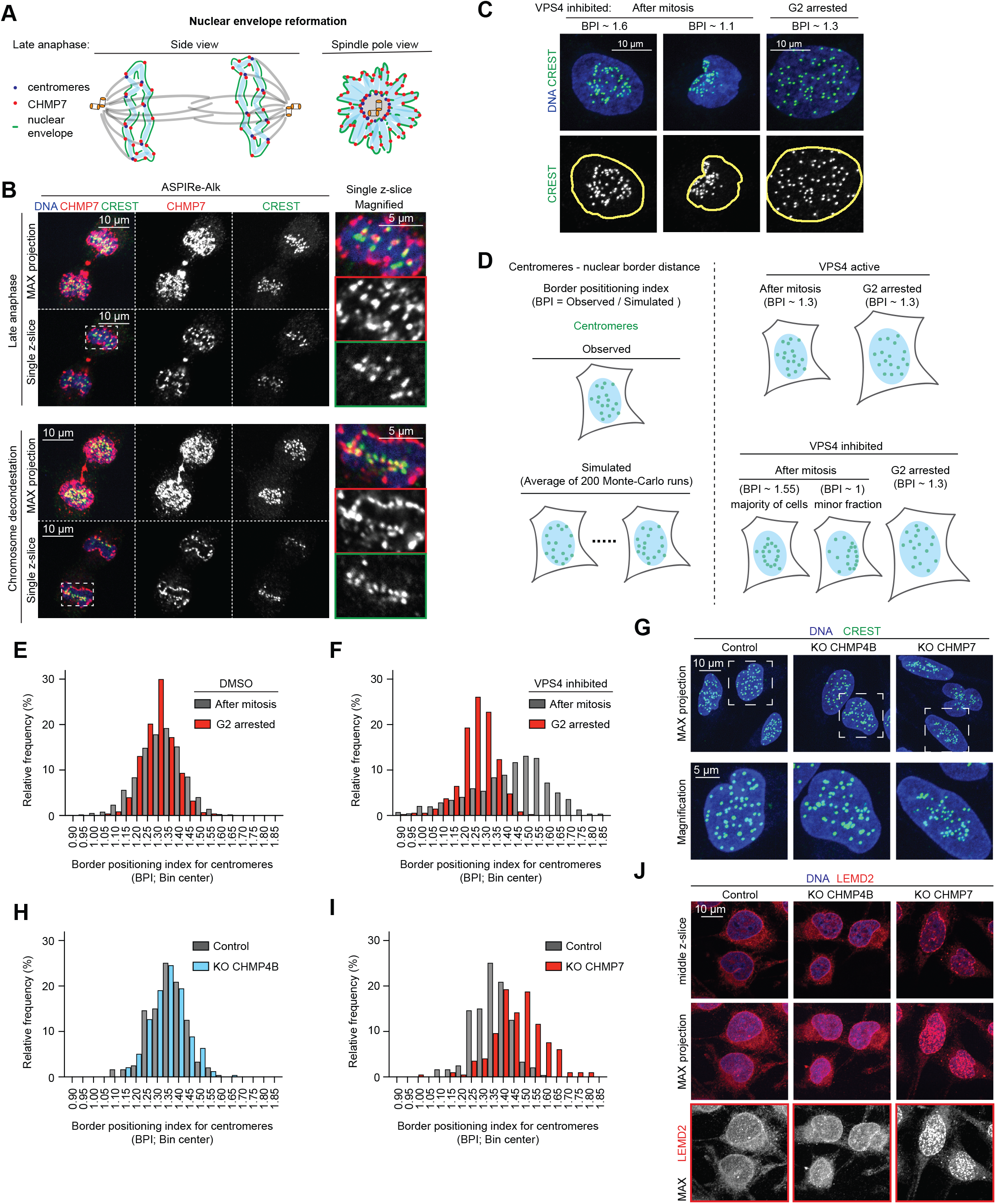
VPS4-mediated turnover of CHMP7 ensures centromere positioning after mitosis. (A) Schematic of CHMP7 and centromere positioning during nuclear envelope reformation. Side and pole views of the mitotic spindle are shown. (B) Representative immunofluorescence images of CHMP7 and centromeres (CREST) following treatment of VPS4B-EC cells with ASPIRe-Alk. Cells were released from a G2 arrest. Maximum intensity projections and single z-slices are shown; scale bars, 10 µm. Magnified boxed region is also shown; scale bar, 5 µm. From top to bottom: merged image, CHMP7 channel, CREST channel. (C) Immunofluorescence images of centromeres following treatment of VPS4B-EC cells with ASPIRe-Alk (5 µM for 18 h). Cells were released from a thymidine block from a thymidine block, and progressed through mitosis or were arrested in G2 for 18 h. The same images as in Figure 4A and 4B are shown. (D) Schematic illustrating the effect of VPS4 inhibition on centromere positioning in post-mitotic nuclei. VPS4 inhibition frequently leads to centromere clustering in ring- or disk-shaped structures at the center of the nucleus. (E) and (F) Analysis of centromere positioning relative to the nuclear border in VPS4B-EC cells following treatment with DMSO or ASPIRe-Alk (5 µM for 18 h). Cells were released from a thymidine block, and progressed through mitosis (gray) were arrested in G2 for 18 h (red). Bars represent the percentage of cells in each bin (bin width, 0.05 BPI); percentages sum to 100% for each condition. Combined data from 3 independent experiments is shown (total number of nuclei for DMSO-treated cells for After mitosis: 1310, G2 arrested: 673; for Alk-treated cells for After mitosis: 1057, G2 arrested: 662) (G) Immunofluorescence images of centromeres in asynchronous Control (parental HeLa), KO CHMP4B, and KO CHMP7 cells. Maximum intensity projections of z-stacks are shown. Scale bar, 10 µm. Magnified boxed regions are also shown; scale bar, 5 µm. (H) and (I) Analysis of centromere positioning relative to the nuclear border in asynchronous Control (parental HeLa), KO CHMP4B, and KO CHMP7 cells. Bars represent the percentage of cells in each bin (bin width, 0.05 BPI); percentages sum to 100% for each condition (total number of nuclei for Control cells: 239; KO CHMP4B: 236; KO CHMP7: 197). The same histogram for the Control is shown in (H) and (I) for comparison. (J) Immunofluorescence images of LEMD2 in Control (parental HeLa), KO CHMP4B, and KO CHMP7 cells. Middle z-slices and maximum intensity projections are shown. Scale bar, 10 µm. See also Figure S5.

To quantitatively analyze centromere positioning relative to the nuclear border, we chose a metric independent of nuclear size or shape. For each nucleus, we calculated the average distance of centromeres to the nuclear border and normalized it to the average from 200 Monte Carlo simulations performed within the same nucleus, to calculate the Border Positioning Index (BPI; Figure 5D and Methods)^62,67^. A BPI > 1 indicates depletion of centromeres from the border relative to a random distribution. We observed a BPI ∼ 1.3 in DMSO-treated cells, both after mitosis and in G2-arrested cells (Figure 5E). VPS4 inhibition shifted centromeres towards the center of the nucleus (BPI ∼ 1.5; Figure 5F). In addition, a subset of cells displayed centromeres positioned closer to the nuclear border (BPI ∼ 1) than in DMSO-treated cells. This repositioning was dependent on mitotic progression and was not observed in G2-arrested cells (Figure 5F).

If VPS4 mediates remodelling of CHMP7-containing complexes to release centromeres from transient associations with the nuclear envelope, then loss of CHMP7 should also lead to the defects in centromere positioning. First, we generated monoclonal knockouts of CHMP7 and CHMP4B (used as a control) in HeLa cells and found both to be viable, although KO CHMP7 cells grew more slowly than control cells (Figure S5C and S5D). Consistent with the model, KO CHMP7 cells displayed centromeres positioned closer to the nuclear center, while KO CHMP4B showed no measurable effect (Figure 5G-5I). Knockout of CHMP7, but not CHMP4B, induced intranuclear LEMD2 and Emerin foci and patchy LEMD2 and Emerin distribution across the nuclear envelope (Figure 5J and S5E). Thus, CHMP7 is required for proper LEM-domain protein localization and centromere positioning, and its loss recapitulates the effects of VPS4 inhibition.

Together, our findings indicate that VPS4/CHMP7 activity is required for proper centromere positioning within the interphase nucleus after mitosis by releasing centromeres from transient interactions with the nuclear envelope during nuclear envelope reformation.

### VPS4 inactivation leads to CHMP7 nuclear foci accumulation followed by DNA damage

We next analyzed VPS4/ESCRT-III functions in the nucleus, focusing on the CHMP7. In untreated cells, besides a transient (∼5 min) recruitment to chromatin during nuclear envelope reformation, CHMP7 displayed prominent accumulation in a fraction of micronuclei, consistent with previous studies^23,58,68^ (Figure S6A). Treatment of VPS4B-EC cells with ASPIRe-Alk resulted in time-dependent accumulation of CHMP7 foci in the nucleus (Figure 6A and 6B). Foci appeared as early as 4 hours after ASPIRe-Alk addition and by 8 hours, the majority of cells displayed the phenotype (>5 spots/nucleus for ∼38 % cells at 4 h, ∼81 % at 8 h and ∼95% at 24 h; average number of spots/nucleus ∼6 in control; ∼36 in Alk-treated; CHMP7 signal per nucleus increased ∼6.1 fold; Figure 6A, 6D, 6E, and S6C). HeLa cells divide every ∼18 h, and the observation that CHMP7 foci first appear in post-mitotic cells is consistent with their appearance during nuclear envelope reformation. Importantly, we observed no effect in Alk-treated VPS4B-WT cells (Figures 6D and S6B), indicating that foci induction is not an off-target effect of the compound. A similar phenotype was also observed in our conditional depletion system (>5 CHMP7 spots/nucleus in ∼96% of cells; average number of spots/nucleus for no Dox: ∼3; +Dox: ∼48; CHMP7 signal per nucleus increased ∼6.3 fold; Figure 6C-6E and S6C).

**Figure 6.**
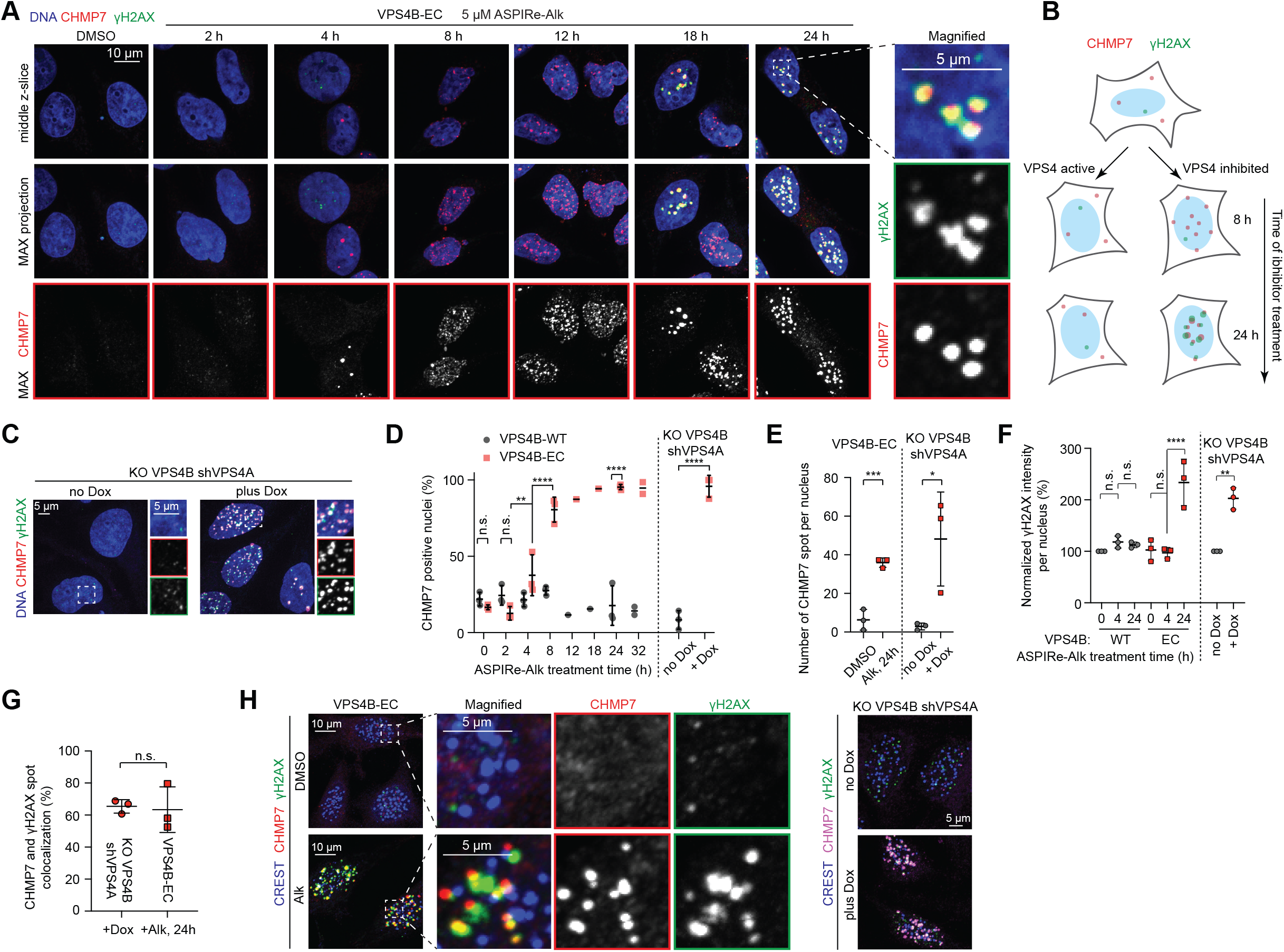
VPS4 inactivation leads to CHMP7 nuclear foci accumulation followed by DNA damage. (A) Representative immunofluorescence images of CHMP7 and γH2AX accumulation in VPS4B-EC cells treated with ASPIRe-Alk (5 µM, indicated time). VPS4 inhibition reveals that nuclear CHMP7 accumulation precedes γH2AX accumulation by ∼12 hours. Middle z-slices and maximum intensity projections are shown; scale bar, 10 µm. Magnified boxed region is also shown; scale bar, 5 µm. (B) Schematic illustrating the effect of VPS4 inhibition on the accumulation of nuclear CHMP7 and γH2AX foci. (C) Immunofluorescence images of CHMP7 and γH2AX accumulation following VPS4 depletion, induced by Dox (2 µg/mL for 48 h). Magnified boxed region is also shown. From top to bottom: merged image, CHMP7 channel, γH2AX channel. Scale bars, 5 µm. (D) Time-dependent accumulation of CHMP7 in the nucleus following VPS4B-EC cells treatment with ASPIRe-Alk (5 µM, indicated time). CHMP7-positive nuclei were defined as nuclei containing >5 spots. Mean values ± SD are shown (points represent independent experiments; >100 cells per data point). One way ANOVA with Šídák’s multiple comparisons test. ****p<0.0001; **p=0.0025; n.s., not significant. For the VPS4 depletion experiment, total cell numbers for no Dox: 435; plus Dox: 275. Unpaired t test; ****p<0.0001. (E) Analysis of the number of CHMP7 spots per nucleus following VPS4B-EC treatment with ASPIRe-Alk (5 µM for 24 h) or upon VPS4 depletion (2 µg/mL for 48 h). The same dataset as in Figure 4D was used for quantification. Mean values ± SD are shown. Unpaired t test; for VPS4 inhibition, ***p=0.001; for VPS4 depletion *p=0.0324. (F) Analysis of γH2AX accumulation following VPS4B-EC treatment with ASPIRe-Alk (5 µM, indicated time) or VPS4 depletion (2 µg/mL of Dox for 48 h). Values were normalized to the DMSO-treated VPS4B-WT condition (100%). Mean values ± SD are shown. A subset of the dataset used in Figure 4D was used for quantification. One way ANOVA with Šídák’s multiple comparisons test. ****p<0.0001; n.s., not significant. For the VPS4 depletion experiment, an unpaired t test was used; **p=0.001. (G) Analysis of the CHMP7 and γH2AX spot colocalization upon VPS4 depletion (2 µg/mL of Dox for 48 h) or following VPS4B-EC treatment with ASPIRe-Alk (5 µM for 24 h). Mean values ± SD are shown (points represent independent experiments); Unpaired t test; n.s., not significant (p=0.8192). (H) Immunofluorescence images of centromeres, CHMP7, and γH2AX in asynchronous cells following VPS4B-EC treatment with Alk (5 µM for 24 h), or VPS4 depletion (2 µg/mL for 48 h). Magnified boxed region is also shown. From left to right: merged image, CHMP7 channel, γH2AX channel. Scale bars are 10 µm and 5 µm, as indicated. See also Figure S6.

The appearance of CHMP7 foci in the nucleus has been linked to DNA damage^23,68^ and we found that VPS4 depletion also results in DNA damage (Figure S1N). Therefore, we examined if the nuclear CHMP7 foci we observed were associated with γH2AX. Interestingly, γH2AX accumulation exhibited a delayed onset relative to CHMP7 appearance and became pronounced at 24 hours of ASPIRe-Alk treatment (∼2.3-fold increase of γH2AX signal/nucleus; Figure 6A and 6F). The majority of γH2AX foci colocalized with CHMP7 (∼63% at 24 h; Figure 6G), and we also observed DNA damage upon VPS4 depletion (∼2-fold increase of γH2AX signal/nucleus; ∼65% of CHMP7 and γH2AX spots colocalized; Figure 6C, 6F, and 6G). Next, we examined if CHMP7 foci, associated with DNA damage, are overlapping with centromeres. In both VPS4 inhibition and VPS4 conditional depletion systems, a fraction of centromeres overlapped with CHMP7-γH2AX signal (Figure 6H).

Together, these findings indicate that VPS4 inactivation first leads to the formation of CHMP7 nuclear foci and subsequently to DNA damage, including damage in the vicinity of centromeres.

### Nuclear CHMP7 foci form independently of other ESCRT-III subunits upon VPS4 inhibition

We observed CHMP7 foci formation in the nucleus (Figure 7A and 6A) and CHMP4B accumulation in the cytoplasm (Figure 7B and 3H-3J) upon VPS4 inhibition, a finding that is inconsistent with models in which these two proteins function together^9,19,47^ (Figure 1D). Therefore, we also examined the localization of other ESCRT-III proteins and VTA1 using immunofluorescence. We found that CHMP1B, CHMP2B, CHMP6, IST1 and VTA1 remain cytoplasmic upon VPS4 inhibition (Figure 7C and S7A-S7D) and obtained similar results in our conditional depletion system. Quantitative analyses of nuclear intensities for each of these proteins was consistent with CHMP7 being the main ESCRT-III protein that forms nuclear foci (Figure 7D and S7E).

**Figure 7.**
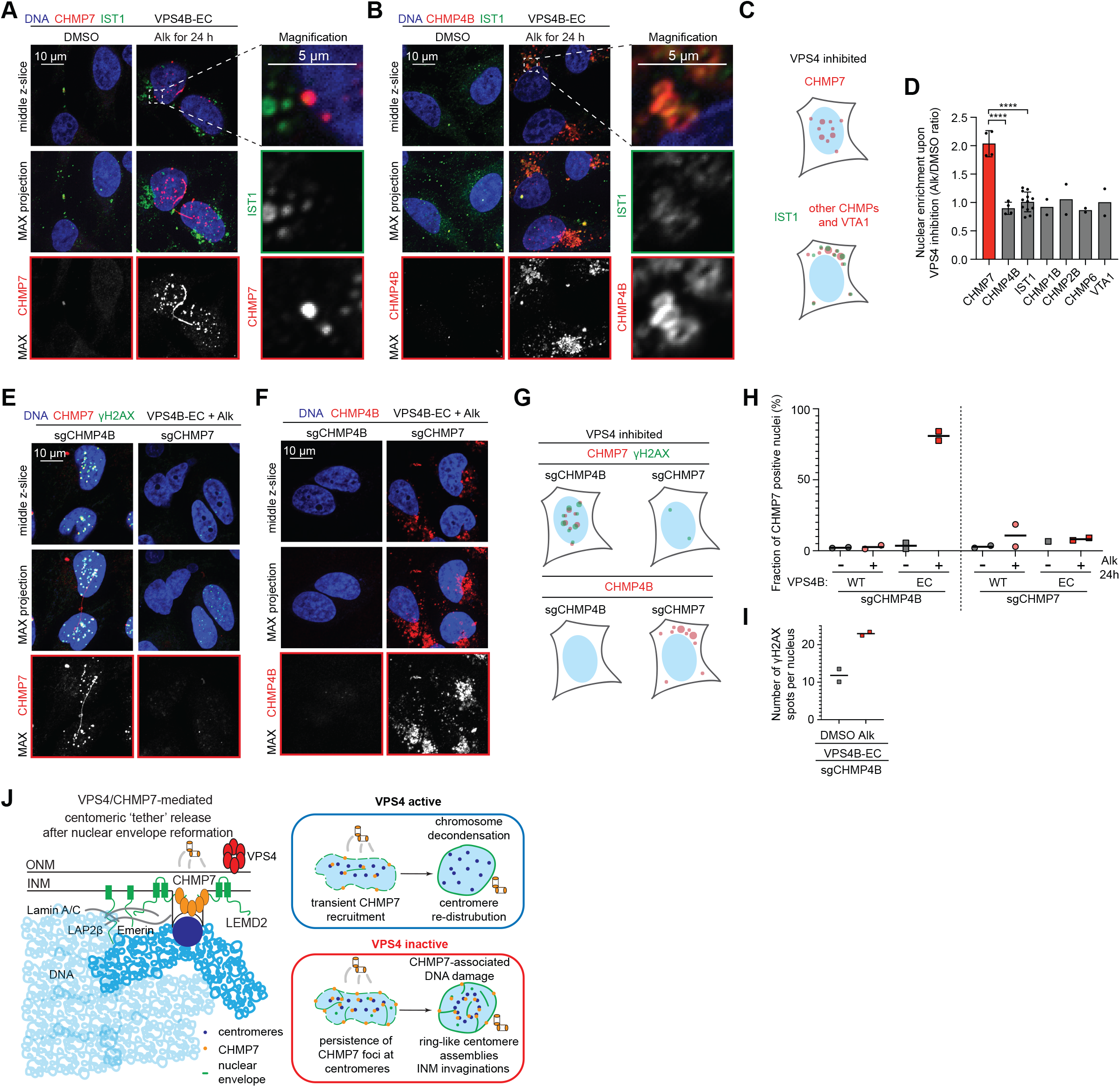
Nuclear CHMP7 foci form independently of other ESCRT-III subunits. (A) and (B) Immunofluorescence images of CHMP7, CHMP4B and IST1 following VPS4B-EC cells treatment with ASPIRe-Alk (5 µM for 24 h). Middle z-slice and maximum intensity projection are shown; scale bar, 10 µm. Magnified boxed region is also shown; scale bar, 5 µm. (C) Schematic of the nuclear and cytoplasmic distribution of ESCRT-III proteins following VPS4 activity inhibition. (D) Analysis of nuclear enrichment of the indicated ESCRT-III proteins following VPS4B-EC cells treatment with ASPIRe-Alk (5 µM for 24 h). Bars are mean values ± SD (points represent independent experiments; >100 cells per data point). ANOVA with Dunnett’s multiple comparisons test; ****p<0.0001. (E) and (F) Immunofluorescence images of CHMP7, γH2AX, and CHMP4B following VPS4B-EC cells treatment with ASPIRe-Alk (5 µM for 24 h). Middle z-slice and maximum intensity projection are shown; scale bar, 10 µm. (G) Schematic of the CHMP7, γH2AX, and CHMP4B accumulation upon VPS4 activity inhibition combined with CHMP4B or CHMP7 depletion. (H) Analysis of CHMP7 accumulation in VPS4B-EC or VPS4B-WT cells depleted of CHMP4B or CHMP7 following ASPIRe-Alk treatment (5 µM, 24 h). (I) Analysis of γH2AX accumulation in CHMP4B-depleted VPS4B-EC cells following ASPIRe-Alk treatment (5 µM, 24 h). (J) Schematic illustrating VPS4/CHMP7-mediated release of centromeres from the nuclear envelope following nuclear envelope reformation. Abbreviations: INM, inner nuclear membrane; ONM, outer nuclear membrane.

Next, depletion of LEMD2, the inner nuclear membrane binding partner of CHMP7, largely prevented CHMP7 nuclear foci formation, consistent with previous studies^19,23^ (Figure S7F). In contrast, depleting individual ESCRT-III subunits, or paralog pairs, or VTA1 neither induced CHMP7 foci in cells with active VPS4, nor prevented their formation upon VPS4 inhibition (Figure S7G-S7J). Thus, CHMP7 foci require the upstream recruiter LEMD2 but not the downstream ESCRT-III subunits, indicating that they are distinct from canonical ESCRT-III polymers.

We then asked whether the DNA damage arising upon VPS4 inhibition shares this dependence. We generated polyclonal CHMP4B and CHMP7 knockouts using CRISPR/Cas9 in cells expressing either the VPS4B-WT or VPS4B-EC allele (Figure 7E and S7K). Upon ASPIRe-Alk-treatment of VPS4B-EC cells depleted of CHMP4B we found that CHMP7 foci accumulated in most nuclei (>5 CHMP7 spots/nucleus in ∼78%; Figure 7E and 7H). In addition, the number of γH2AX foci also increased (spots/nucleus for control: 12, treated: 23; Figure 7I). Co-localization analysis showed that ∼49% of CHMP7 spots overlapped with γH2AX signals (Figure 7E). As would be expected, we did not observe nuclear CHMP7 foci formation upon VPS4B inhibition in cells depleted of CHMP7 (Figure 7E and 7H). Importantly, CHMP7 depletion also suppressed the appearance of γH2AX foci, indicating reduced DNA damage (Figure 7E and 7G). In contrast, CHMP7 depletion did not prevent the accumulation of CHMP4B foci in the cytoplasm (Figure 7F and 7G), and depletion of other ESCRT-III subunits did not prevent the appearance of CHMP7-associated γH2AX (Figure S7G-S7J).

Together, these findings indicate that the nuclear CHMP7 foci formed upon VPS4 inactivation are distinct from canonical ESCRT-III polymers and that the resulting DNA damage depends specifically on CHMP7, not on CHMP4B or other ESCRT-III subunits. This damage in part arises near the mispositioned centromeres, linking the persistence of CHMP7 to the loss of genome integrity.

## Discussion

Our findings reveal that centromere positioning in the nucleus following mitosis is a regulated step that depends on VPS4/CHMP7. Upon acute chemical inhibition of VPS4, CHMP7 accumulates in nuclear foci and persists at a subset of centromeres, which remain constrained at sites established by the mitotic spindle. This improper CHMP7 localization is accompanied by defects in nuclear envelope organization, altered distribution of nuclear envelope proteins and formation of inner nuclear membrane invaginations. Over time, DNA damage accumulates in the vicinity of centromeres. Nuclear CHMP7 foci are largely devoid of other ESCRT-III subunits and CHMP7 depletion, but not the loss of CHMP4B or other ESCRT-III subunits, suppresses DNA damage upon VPS4 inhibition. These results support a model in which VPS4-mediated turnover of CHMP7 is required to release centromeres from transient nuclear envelope associations, enabling their proper positioning in the daughter nuclei after each mitosis.

Based on our data we propose a model for how VPS4 and CHMP7 ensure proper post-mitotic centromere positioning. During prometaphase, chromosomes gather at the cell equator with their arms oriented outward and centromeres clustered in a ring-like configuration, resulting from the spindle shape of the cell division apparatus^63^. As cells progress into anaphase, chromosomes move toward the spindle poles through shortening of kinetochore microtubules, such that centromere positions at the end of anaphase represent projections of their metaphase arrangement along the spindle axis. Viewed from the pole, this organization appears as a ring- like arrangement of centromeres, in part due to exclusion of chromosome arms from the central spindle by interpolar microtubules^63^. Under normal conditions, this geometry is transient and resolves following nuclear envelope reformation and chromatin decondensation, allowing centromeres to redistribute in the interphase nucleus. Consistent with this timing, our imaging indicates that VPS4 and ESCRT-III subunits are recruited to chromatin ∼10 minutes after anaphase onset, coinciding with LEMD2 recruitment and restoration of nuclear import. CHMP7 and other ESCRT-III subunits dissociate from chromatin within ∼5 minutes of recruitment, with centromeres representing the final sites of detectable CHMP7 association. VPS4 inactivation prevents clearance of CHMP7 and LEM-domain proteins from centromeres, locking them in persistent association with each other^19,21,26^. We propose that the nuclear membrane invaginations observed under these conditions do not create detectable gaps in the nuclear envelope but instead remain stably linked to centromeres. The persistence of these invaginations, likely tethered to centromeres through LEM-domain protein/Lamin A/C-containing complexes, prevents centromere redistribution after mitosis (Figure 7J).

Current models for ESCRT-III function during nuclear envelope reformation suggest that CHMP4B and other ESCRT-III subunits are recruited downstream of the LEMD2-CHMP7 complex to mediate membrane sealing^9,19,47^. Our observations from live-cell microscopy and immunofluorescence experiments under control conditions are consistent with these models. However, our findings suggest that CHMP7 assemblies formed upon VPS4 inactivation are distinct from canonical ESCRT-III polymers. These nuclear CHMP7 foci are largely devoid of other ESCRT-III subunits, and while both CHMP7 and LEMD2 form nuclear foci upon VPS4 inhibition, LEMD2 also accumulates in the cytoplasm under these conditions. Notably, the formation of CHMP7 foci depends on LEMD2 but does not require other ESCRT-III proteins. One possible explanation is that, during mitosis, nuclear and cytoplasmic contents mix, and the subsequent nuclear accumulation of CHMP7 reflects its retention at reforming nuclear membranes and its indirect association with centromeres following mitotic exit. The absence of other ESCRT-III subunits at CHMP7 foci may result from their preferential recruitment to cytoplasmic assembly sites, suggesting competition for ESCRT-III proteins between distinct cellular locations. Indeed, upon VPS4 inhibition, we observed accumulation of ESCRT-III proteins in the perinuclear region near the Golgi apparatus, consistent with impaired endosomal maturation.

A result we did not anticipate was the dispensability of CHMP7, the only known recruiter of ESCRT-III to the nuclear envelope^9,19,47^. That nuclear envelope sealing can proceed in CHMP7-depleted HeLa cells implies the existence of compensatory mechanisms and suggests that ESCRT-III subunit requirements vary by cell type. Previous studies indicated a delayed but not fully abrogated repair of nuclear envelope ruptures upon ESCRT-III subunit depletion in migrating cancer cells^69,70^. ESCRT-III independent mechanisms of nuclear envelope sealing mechanisms have been identified in fission yeast, *S. japonicus*^71,72^. These mechanisms include regulation of membrane lipid composition through the fatty acid desaturase Ole1 and the recently reported Alx1/Vid27 pathway, in which Vid27 localizes to sites of post-mitotic nuclear envelope sealing and functions downstream of the ESCRT-III adaptor Alx1^72^. However, no Vid27 ortholog has thus far been identified in humans. In the related fission yeast *S.pombe*, spindle pole body components have also been implicated in preserving nuclear compartmentalization^73^. The observation of CHMP7 dispensability extends to other ESCRT-III components: individual subunit depletions, including combined disruption of both paralogs CHMP1A/B and CHMP2A/B, are compatible with HeLa cell growth, contrasting with the essentiality of VPS4 across ESCRT-III-dependent processes. DepMap data corroborate this variability^46^, revealing divergent phenotypic consequences of CHMP7, CHMP4B, and CHMP2A depletions across cell lines, with approximately half of ESCRT-III subunits appearing broadly dispensable. Our findings suggest that ESCRT-III polymers have variable subunit composition, and that individual subunits may serve both redundant and specialized functions in a cell type-specific manner.

Acute inhibition of VPS4 reveals a ∼12-hour delay between the appearance of nuclear CHMP7 foci and the onset of DNA damage, consistent with a model in which DNA damage arises as a secondary consequence of unresolved CHMP7/LEM-domain protein assemblies at the inner nuclear membrane. This delay may reflect progressive CHMP7 polymerization-induced torsional stress on DNA, associated with the inner nuclear membrane, resembling previously observed phenotypes upon overexpression of a nuclear export-deficient CHMP7 mutant^23^. Alternatively, and not mutually exclusively, a fraction of the observed DNA damage may arise from local rupture of the inner nuclear membrane following prolonged deformation by CHMP7 assemblies. However, such rupture was not observed within the limits of our EM analysis and is not supported by preserved cytoplasmic compartmentalization of ESCRT-III components other than CHMP7. The rescue of DNA damage by CHMP7 depletion, but not by depletion of other ESCRT-III proteins, highlights the deleterious effects of unresolved CHMP7 polymers as part of the VPS4 inhibition phenotype and reveals unique function of CHMP7.

The functional relevance of VPS4/CHMP7-mediated nuclear envelope remodeling is not restricted to dividing cells. In cortical neurons, VPS4/ESCRT-III has been proposed to regulate nuclear envelope remodeling through a complex involving the DNA-binding scaffold protein SATB2 and LEMD2^74,75^. Our findings from VPS4 inhibition in post-mitotic and G2-arrested cells, together with CHMP7 knockout experiments, support a model in which VPS4/CHMP7 acts as a remodeler of chromatin-nuclear envelope contacts in human cells, illustrated by the ring-like positioning of centromeres in post-mitotic nuclei. This function is conceptually analogous to the role established for ESCRT-III/Vps4 in *S. japonicus*^26^ and is consistent with the centromere-tethering and pericentromeric heterochromatin-silencing functions attributed to Lem2 in the related fission yeast *S. pombe*^76–78^. Prior evidence for a nuclear envelope-centromere connection in human cells also comes from the progeria-associated Lamin A E145K mutation, which induces aberrant centromere clustering in the nuclear interior^64,79^. This phenotype has been reported to arise during nuclear envelope reassembly and requires passage through mitosis. Biochemical interactions among Emerin, Lamin A, BAF, and centromeric proteins suggest a possible molecular basis for this connection^64^. Further supporting a role for VPS4/CHMP7/ESCRT-III in inner nuclear membrane remodeling beyond post-mitotic nuclear envelope sealing, depletion of the ESCRT-III subunit CHMP4B has been reported to induce aberrant inner nuclear membrane invaginations in human cells^80^. Together with the identification of LBR and LEM-domain proteins as determinants of peripheral centromere positioning^27^, our findings suggest that VPS4/ESCRT-III functions as a disassembler of LEM-domain protein/Lamin A chromatin tethers in human cells.

Our chemical genetics analyses address two key gaps in the evaluation of VPS4 paralogs as a therapeutically actionable vulnerability in cancer^29,39^. First, we show that inhibition of VPS4 catalytic activity, rather than loss of the protein, is sufficient to kill cells lacking one VPS4 paralog, but is not toxic in cells expressing both. Second, VPS4 is believed to function as a hexamer to remodel and disassemble ESCRT-III polymers^81^, and both VPS4A and VPS4B may co-assemble into mixed hexamers in cells expressing both paralogs. Therefore, it is possible that inhibiting one paralog would be toxic to normal cells in which mixed hexamers are present and functional. Our finding that allele-specific inhibition of VPS4B is not toxic to cells that express wildtype VPS4A (Figure S2C) argues against this possibility. Further studies across different cell types will be needed to fully address this question, particularly as paralog-selective VPS4 inhibitors have not been reported and their design is likely to be challenging due to the high sequence similarity between the two paralogs.

The chemical genetics approach we have developed could be used to selectively and acutely inhibit VPS4 function in the context of other dynamic cellular processes that involve membrane remodeling, ranging from HIV egress^82,83^ to the export of large transcripts in muscle cells^84^. This approach could also be extended to other AAA proteins for which selective inhibitors have not been developed and would allow dissecting fundamental cellular mechanisms and testing therapeutic hypotheses.

## Data and resource availability

Source Data contains uncropped western blots and data used to build graphs presented in figures. Compound synthesis is described in Supplementary Information. The code used for analysis of spatial centromere distribution is available on GitHub (link). Key plasmids have been deposited to Addgene. Every effort will be made to share data and materials, assuming funds are available to cover costs.

## Declaration of generative AI and AI-assisted technologies in the writing process

During the preparation of this work, the authors used ChatGPT (OpenAI) and Claude (Anthropic) to analyze grammar and correct typos. After using these tools, the authors reviewed and edited the content as necessary and take full responsibility for the content of the published article.

## Acknowledgements

This work was funded by a National Institutes of Health grant to T.M.K. (R35GM130234); N.K. was supported by this NIH award, and in part by funds from a Merck Postdoctoral Fellowship at The Rockefeller University; T.H. is grateful to the NIH/NCI (F32CA284727) for support. The authors thank Professors Katya Vinogradova and Hiro Funabiki at The Rockefeller University for generously sharing reagents and equipment. We appreciate discussions with Amol Aher (currently at the ETH, Zurich). The authors thank Yang Xiao and Leishemba Thoidingjam for their support in compound synthesis. We are also grateful to Anurag Sharma at the Electron Microscopy Resource Center at The Rockefeller University.

## Author Contributions

N.K. conducted and analyzed the majority of experiments, wrote the code for image analysis and prepared figures. T.H. and N.J. designed and synthesized compounds. T.H. and A.K. performed protein biochemistry. M.K. contributed to live-cell imaging experiments. R.T. contributed to microscopy data acquisition and quantifications. H.A.P. processed electron microscopy samples. N.K and T.M.K. designed the study and wrote the manuscript with input from all authors.

### The authors declare no competing interests

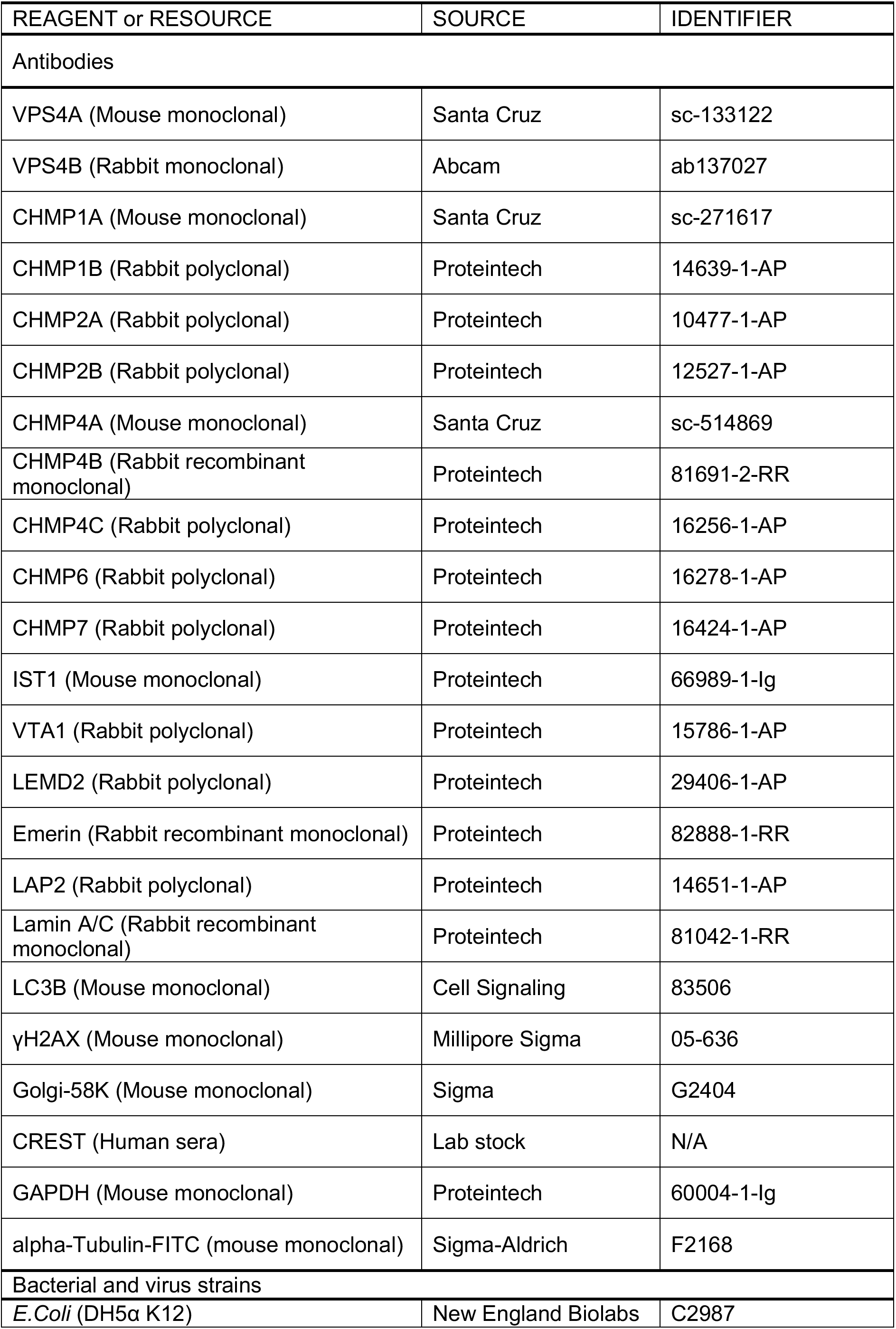
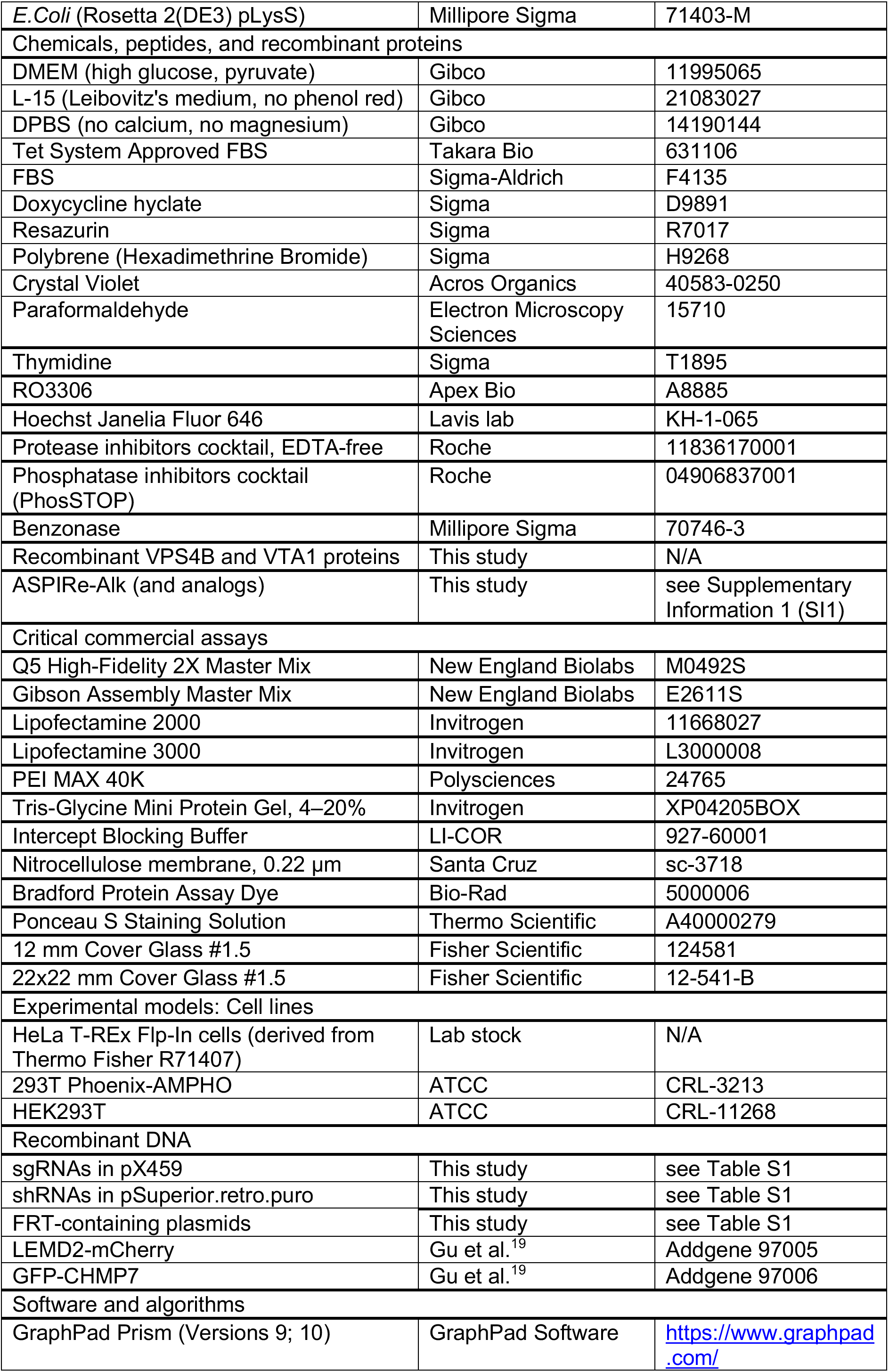
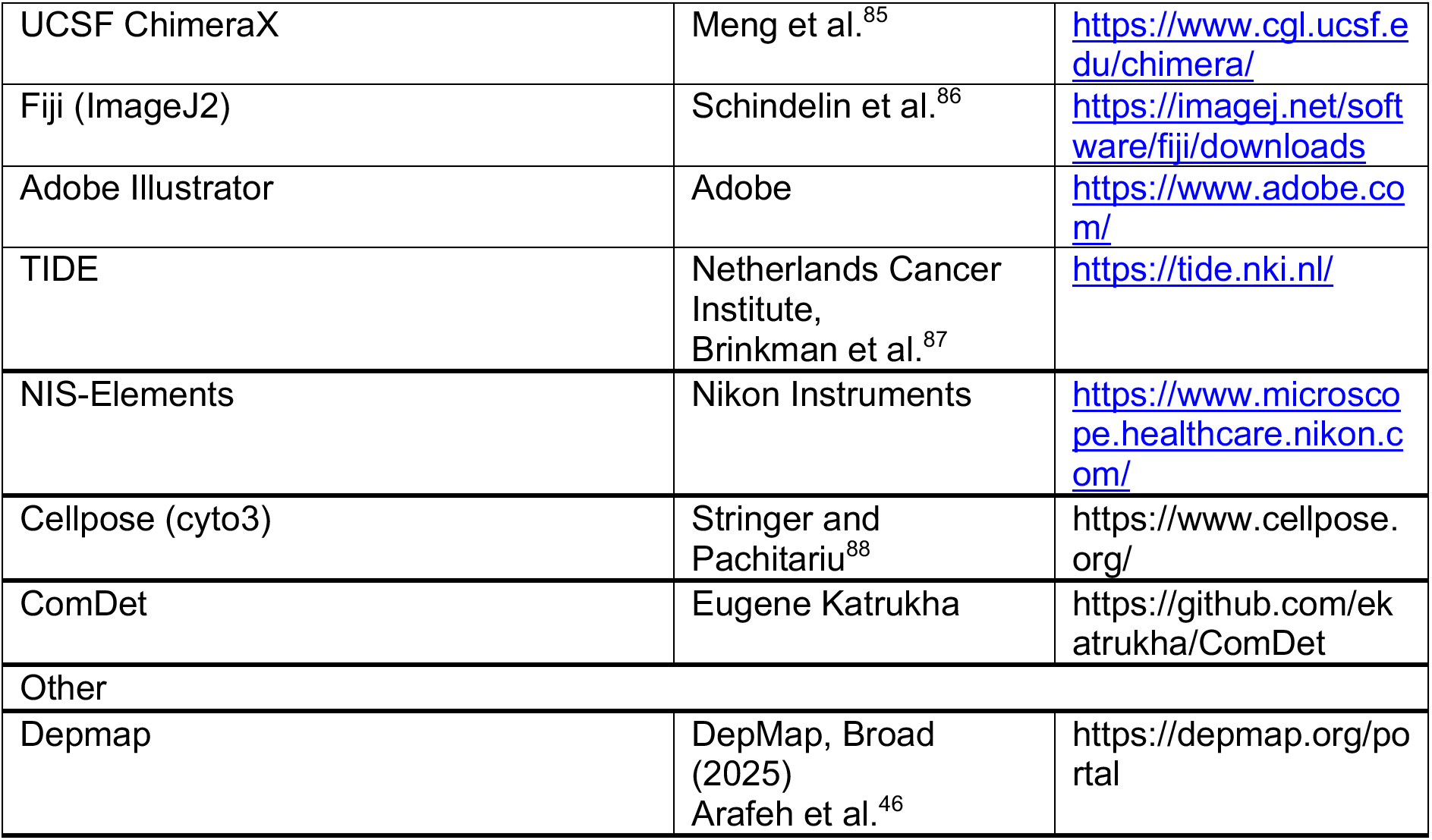
Key resources table.

### Experimental models

HeLa T-REx Flp-In cells (HeLa; derived from Thermo Fisher R71407) from the lab stock were used as a parental cell line for cells presented in this study, unless otherwise mentioned. Cells were cultured in high-glucose DMEM with sodium pyruvate and glutamine (Gibco 11995065) supplemented with 10% FBS (Sigma-Aldrich F4135), without antibiotics. Cell lines expressing doxycycline (Dox)-inducible constructs were generated and maintained in DMEM supplemented with 10% Tet System Approved FBS (Takara Bio 631106). For the live-cell imaging cells were maintained in Leibovitz’s L-15 media (no phenol red; Gibco 21083027) supplemented with 10% FBS. Key cell lines were tested for mycoplasma and confirmed to be negative before and after genetic engineering. Testing was done using a PCR-based method^89^. Cells were cultured in a humidified incubator at 37°C and 5% CO2.

## Method details

### Transfection

Cells were transfected using Lipofectamine 2000 (Invitrogen 11668027), Lipofectamine 3000 (Invitrogen L3000008), or PEI MAX 40K (PEI; Polysciences 24765) according to manufacturer’s instructions. A brief protocol for PEI transfection of HeLa cells (for a well of 6-well plate) provided below. Actively growing cells were seeded to achieve ∼80% confluency at the time of transfection. Typically, ∼40×10^4^ cells/well were plated in the evening, and transfection was performed 24 h later (∼100×10^4^ cells). A 4-fold mass excess of PEI relative to DNA was used. A total of 2.5 μg of DNA and 10 μg of PEI were diluted separately in Opti-MEM, combined and mixed by tapping, and incubated at room temperature for 30 min. The mix was then added dropwise to the cells. The next morning, the medium was replaced with fresh.

### Flp-In cell line generation

FRT-containing plasmids were co-transfected with the flippase encoding pOG44 plasmid at a 1:9 mass ratio (e.g., 250 ng FRT plasmid and 2250 ng pOG44). 48 hours after transfection, cells were reseeded into larger culture dishes in medium supplemented with 250 μg/mL of Hygromycin B. Medium was replaced every three days, with a DPBS wash to remove dead cells. Seven days after the start of selection, well-visible resistant colonies were observed in transfected cells, while no cells remained in the no-plasmid control. After 10 days of selection, cells were detached and reseeded into smaller dishes while maintaining selection. Total antibiotic selection lasted 12-14 days. After selection, cells were passaged once without antibiotic, and transgene integration was confirmed by methods of choice, including immunoblotting, PCR from genomic DNA, or fluorescence microscopy.

### Immunoblotting

For immunoblotting, cells were washed once with DPBS (Gibco 14190144) and either lysed directly on the plate or trypsinized, neutralized with complete medium, washed with DPBS, flash-frozen and stored at -80°C until further use. Lysis was made in RIPA buffer (50 mM Tris-HCl, pH 8.0, 150 mM NaCl, 0.1% Triton X-100, 0.5% sodium deoxycholate, 0.1% SDS) supplemented with 0.1 mM PMSF, protease and phosphatase inhibitors, 3 mM MgCl2, and ∼25 U/mL of Benzonase immediately before use. Lysis was conducted for 30 min at 4°C with pipetting, and the non-clarified lysate was collected. Protein concentrations were measured using a Bradford assay (Bio-Rad 5000006) and samples were mixed with 5x Laemmli loading buffer (50 mM Tris-HCl, 10% SDS, 5% 2-mercaptoethanol, 10% glycerol, 0.01% bromophenol blue; pH adjusted to 6.8). Samples were denatured at 95°C for 5 min, cooled to room temperature, and centrifuged at 14000 g for 5 min at room temperature. Samples were resolved by SDS-PAGE using either precast gels (gradient 4-20%; Invitrogen XP04205BOX) or self-cast gels (10% or 12%). Proteins were transferred to nitrocellulose membrane (Santa Cruz sc-3718) by wet transfer 40 V overnight at 4°C. To check transfer and loading, membranes were stained with Ponceau Red (Thermo Scientific A40000279), then blocked for 1 h at room temperature using Intercept Blocking Buffer (LI-COR 927-60001). After blocking, membranes were incubated with primary antibodies diluted in blocking buffer for 2 h at room temperature or overnight at 4°C, followed by 3×10 min washes in TBST. Membranes were incubated with secondary antibodies for 1 h at room temperature, washed 4×5 min in TBST, and imaged using an Odyssey scanner (LI-COR) or GelDoc imager (Bio-Rad). Antibody dilutions are listed in Table S2.

### Immunofluorescence

Actively growing cells were seeded on sterile uncoated or poly-L-lysine coated 12 mm round coverslips #1.5 (Fisher Scientific 124581). Cells were allowed to attach and grow for 24 h prior to the start of experiments. For initial experiments, two fixation protocols were used: fixation with ice-cold methanol for 10 min at -20°C, or fixation with 3.2% paraformaldehyde in DPBS for 15 min at room temperature. After fixation, coverslips were washed 3×5 min in DPBS. For paraformaldehyde fixation, the second wash contained 125 mM glycine. Further coverslips were incubated in a blocking buffer (3% BSA, 0.2% Triton X-100 in DPBS) for 1 h at room temperature. Coverslips were incubated with primary antibodies diluted in blocking buffer either overnight at 4°C or 2 h at room temperature, followed by 3×5 min wash in DPBS. Secondary antibodies incubation was combined with DNA staining (500 nM; Hoechst JF646) for 1 h at room temperature, followed by 3×5 min washes in DPBS. Coverslips were put in drops of mounting medium (20 mM Tris pH 8.0, 0.5% n-propyl gallate, 90% glycerol) on Superfrost slides (Fisher Scientific 1255015), allowed to dry in the dark, and excess mounting medium was removed by aspiration. Coverslips were sealed with transparent nail polish, and slides were stored at -20°C before image acquisition. As both fixation methods showed similar results in initial tests, methanol fixation was used further for all data present in the figures. Antibody dilutions are listed in Table S2.

### Plasmids

All plasmids used in this study were generated using standard molecular biology techniques by PCR amplification from plasmid templates (New England Biolabs; Q5 High-Fidelity 2X Master Mix M0492S), followed by Gibson assembly (Gibson Assembly Master Mix; New England Biolabs E2611S) into destination vectors. Key plasmids are listed in Table S1.

### Image acquisition

Images were acquired using NIS-Elements software on a Nikon Ti2-E microscope equipped with a W1 confocal scanning unit and a custom Yokogawa quad notch filter (405/480/561/640 nm) for excitation. Emission light was filtered using either ET 520/40m, ET 615/40m, or ET 685/50m filters (Chroma Technology). Typical imaging parameters for immunofluorescence were: 488 nm (100 mW laser, 40% power, 100 ms exposure); 561 nm (80 mW laser, 40% power, 100 ms exposure; 640 nm 75 mW laser, 40% power, 100 ms exposure). A 100x 1.45 NA Plan Apo objective (Nikon) and a Photometrics Prime 95B camera were used. 35 slices with a 0.3 μm step were acquired. For larger fields of view, 3×3 image scans tiled images with 15% overlap were acquired and stitched using blending. Acquired images had a resolution 3240 x 3240 pixels (356.4 x 356.4 µm).

### Live-cell imaging

For live-cell imaging, cells were seeded on 22×22 mm #1.5 coverslips (Fisher Scientific 12-541-B) 48 h prior to imaging. To induce expression of mNG-tagged constructs under Tet-inducible promoters, 100 ng/ml doxycycline was added 24 h before imaging. To visualize DNA, 125 nM of Hoechst JF646 at final 0.1% DMSO was added 1 h before imaging and maintained during the experiment. Coverslips were assembled in modified Rose chambers containing Leibovitz’s L-15 media (no phenol red; Gibco 21083027), supplemented with 10% FBS, DNA stain, and doxycycline. Metaphase cells were identified and imaged further at 1 min intervals. Number of z-slices for a width (z-stacks were acquired with an appropriate number of slices to cover the full cell volume).

### Electron Microscopy

VPS4B-EC cells were seeded into a 6-well plate and allowed to attach and grow for 24 h before treatment with DMSO or 5 μM ASPIRe-Alk for 18 h. Cells were fixed in 2% glutaraldehyde in 0.1 M sodium cacodylate buffer (pH 7.2) 2 mM CaCl2 for 4 hs at room temperature followed by overnight fixation at 4C, postfixed in 1% osmium/0.8% potassium ferricyanide in 0.1 M cacodylate buffer, followed by post-staining in 1% uranyl acetate in 0.05 M maleate buffer pH 5.2, dehydration in an ethanol series, and embedding in Eponate 12 (Ted Pella, Inc). Ultrathin sections (60-65 nm) were stained with uranyl acetate and lead citrate, and images were acquired using a Tecnai G2-Spirit transmission electron microscope (FEI, Hillsboro, Oregon) operated at 120 kV, equipped with an AMT BioSprint29 digital camera (AMT, Danvers, MA).

### CRISPR-mediated knockouts

sgRNAs targeting gene-of-interest were designed using CHOPCHOP (https://chopchop.cbu.uib.no/)^90^ or CRISPick^91^, and in general followed G(N)_19_ rule. For the negative control sgRNA targeting gene desert on chromosome 2 was used (sgContol), similar to previously published^29^. As a control for pan-essential gene, housekeeping gene VCP (p97) was used. Targeting of KIF11 (kinesin-5) was used as a control for gene required for mitotic progression. sgRNAs were inserted into pX459 all-in-one plasmid (Addgene_62988)^92^ by ligation of annealed oligonucleotides (IDT DNA). Plasmids were verified by sequencing (Azenta) and sterile DNA that was used for cell transfection was prepared using the Qiagen midiprep kit. 48 h after transfection HeLa cells were reseeded in medium containing 2 µg/ml puromycin for 24 h. Puromycin was washed away by 2x DPBS washes and fresh medium was added for 48 h. Knockout efficiency was addressed using immunoblotting, efficient sgRNAs were identified and used further. Monoclonal knockout cell lines were prepared by single clone isolation by limited dilution. Briefly, cells were diluted to 0.6 cells/well and seeded into the clear-bottom 96 well plates. 6 days after seeding wells containing single growing clones were identified by light microscopy examination and labelled, media was replaced for fresh. 10 days after seeding, colonies were detached and dispersed, and cells reseeded into the 12 well plates, where they were growing with medium exchange until subconfluency was reached. Part of the cells were used for immunoblotting to test for loss of the gene expression and part was used for genomic DNA isolation (DNeasy Blood and Tissue kit, Qiagen), followed by PCR amplification of a region containing a cut site. Amplicons were Sanger sequenced (Azenta) and analyzed for the knockout efficiency using TIDE^87^. For final preparation of the knockout cell lines, three verified clones were mixed together to reduce effects of clonal variation. sgRNA sequences are listed in Table S1.

### shRNA-mediated VPS4 depletion

To deplete VPS4 paralogs, selected shRNAs were cloned into the pSuperior.retro.puro vector. The vector was transfected into HEK293T Phoenix-AMPHO cells (ATCC CRL-3213) to generate virus-like particles. 48 h after transfection, culture medium containing virus-like particles was collected and filtered through a 0.45 µm filter to remove cell debris. The medium was supplemented with 4 µg/ml polybrene and added to HeLa cells. 48 h after transduction, HeLa cells were reseeded in medium containing 2 µg/ml puromycin for 72 h. Cells were passaged once without antibiotic before testing depletion. For depletion experiments cells were seeded at ∼20% confluency, allowed to attach and grow for 24 h. Depletion was induced by addition of 2 µg/ml doxycycline for 48 h. To prepare the final KO VPS4B shVPS4A cell line, three verified individual clones demonstrating high depletion efficiency were mixed together, to minimize effects of clonal variation. shRNA sequences are listed in Table S1.

### 96-well plate based cell viability assays

Cells were seeded into the 96 well plates at 10^4^ cells/well and allowed to attach for 2-4 hours. Then assayed compounds were added to the desired concentrations: doxycycline for shRNA-mediated depletion or ASPIRe-Alk/ASPIRe-analogs for inhibition of VPS4B cysteine alleles. For ASPIR-analogs, medium contained final 0.1% DMSO. Further cells were growing for 72 h and cell viability was measured by Resazurin assay. Final 0.05 mM Resazurin (Sigma R7017) was added to the wells at 70 h of growth for 2 h, followed by fluorescence measurement using plate reader (excitation 550 nm, emission 590 nm; Synergy NEO Microplate Reader). Fluorescence measurements were background corrected (wells with no cells) and normalized to corresponding unperturbed controls, taken as a 100% value. Data was analyzed in GraphPad Prism using sigmoidal least squares fit ([Inhibitor] vs. normalized response --Variable slope), described by Y=100/(1+(IC_50_/X)^HillSlope).

### Colony formation and cell viability assays

For colony formation assays, actively growing cells were seeded at 500 or 1000 cells/well of 24-well plate. Cells were allowed to attach and divide for 24 h. Then experimental conditions were applied by medium exchange. Medium was refreshed every 3 days, keeping experimental conditions during the 7-9 days of assay. At 7 days of growth ∼1 mm size cell colonies were visible in control wells. Further, cells were washed once with DPBS and fixed with methanol for 15 min at room temperature. Cells were stained with 0.5% Crystal Violet in 25% methanol for 30 min at room temperature and then washed with water 4×5 min with shaking, until dye stopped to be released. Plates were air-dried overnight in the dark and scanned using Odyssey scanner (LI-COR). For the cell viability assays equal amounts of cells between different conditions were plated for indicated time, typically 48 h or 72 h, and further fixed and stained similar to colony formation assays.

### Cell synchronization and release

To synchronize HeLa cells in early S phase, they were treated with 2 mM thymidine for 18 h. G2 arrest was achieved by CDK1 inhibition using 10 μM RO3306, treated with indicated times, up to 18 h. Release was achieved by washing away treatment agents with two washes of pre-warmed DPBS followed by wash with complete medium, with careful shaking ∼30 sec during each wash.

### Click chemistry reactions

Cells were treated with ASPIRe-Alk, washed with PBS, and lysed in RIPA buffer supplemented with protease inhibitors, PMSF, phosphatase inhibitors, benzonase, and MgCl₂, as described above. Protein concentrations were normalized, and the click reaction was performed by adding THPTA (final 200 µM), CuSO₄ (2 mM), azide rhodamine (50 µM), and TCEP (1 mM). The reactions were incubated for 30 min at 37 °C with shaking while protected from light and were then quenched by EDTA followed by adding Laemmli loading dye and boiling 5 min at 95 °C. Proteins were analyzed by SDS-PAGE, followed by in-gel fluorescence imaging and immunoblotting.

### Multiple sequence alignment

Protein sequences were derived from KEGG database and organismal codes are given according to KEGG nomenclature^93^ (https://www.genome.jp/kegg/). Multiple sequence alignments were prepared in Unipro UGENE^94^ (https://ugene.net/) using Clustal Omega, with gray color intensity indicating percentage of identity. Selected alignment regions were exported, and the final look was adjusted in Adobe Illustrator.

### Recombinant protein purification

Protein expression and purification was done similar to what was previously described^37^. Below is a typical protein expression and purification protocol for His6Sumo-VPS4B constructs and GST-VTA1, human protein sequences. Proteins were expressed in *E.Coli* (Rosetta 2(DE3) pLysS; Millipore Sigma 71403-M) grown in Miller’s LB medium. Expression was induced with 1.0 mM IPTG (Goldbio) at OD600 ∼0.6-0.8. Cultures were grown with vigorous shaking at 18°C for 12-16 hours, pelleted, and resuspended in a lysis buffer (for VPSB constructs: 20 mM HEPES-KOH, 500 mM KCl, 4 mM MgCl₂, 20 mM Imidazole, 5% Glycerol, 10 mM 2-mercaptoethanol, 1 mM ATP, pH 7.6; for VTA1: 20 mM HEPES-KOH, 250 mM KCl, 2 mM MgCl₂, 5% Glycerol, 2 mM DTT, 0.2 mM ATP, pH 7.6) supplemented with benzonase, 1 mM PMSF, and protease inhibitor cocktail at 4°C. Cells were lysed using Emulsiflex-C5 homogenizer (Avestin, 5-6 cycles at 10,000-15,000 psi). The homogenized lysate was clarified by centrifugation at 100,000g for 45-60 min using either a Ti70 or Ti45 rotor in a Beckman Coulter Optima LE-80K ultracentrifuge; lysates were filtered and proteins were purified by following way. VPS4B constructs were purified using metal-affinity chromatography using 5 mL HisTrap HP columns, lysates were loaded on the column followed up by extensive column wash with the lysis buffer (10 column volumes) and eluted in 20 mM HEPES-KOH, 500 mM KCl, 2 mM MgCl₂, 400 mM Imidazole, 5% Glycerol, 10 mM 2-mercaptoethanol, pH 7.6. His6-Sumo tag was cleaved by incubation with His6-Ulp1 protease combined with dialysis into 20 mM HEPES-KOH, 50 mM KCl, 20 mM Imidazole, 2 mM MgCl₂, 5% Glycerol, 10 mM 2-mercaptoethanol, pH 7.6 overnight at 4°C. To capture cleaved His6-SUMO tag and His6-Ulp1 protease, dialysed proteins were incubated with Ni-NTA resin for 20 min. Flowthrough was collected and proceeded for ion-exchange chromatography. Proteins were loaded on a MonoQ 5/50 GL column (Cytiva) pre-equilibrated with a low salt buffer (20 mM HEPES-KOH, 50 mM KCl, 2 mM MgCl₂, 5% Glycerol, 2 mM DTT, pH 7.6). The column was washed with 3 column volumes of 3% high salt buffer (20 mM HEPES-KOH, 1 M KCl, 2 mM MgCl₂, 5% Glycerol, 2 mM DTT, pH 7.6) and eluted in a 3%-25% gradient of high salt buffer over 25 column volumes. Fractions containing VPS4B were concentrated using an Amicon Ultra 30K concentrator, 0.22 μm filtered, and loaded on a Superdex 200 Increase 10/300 GL column (Cytiva) equilibrated in 20 mM HEPES-KOH, 150 mM KCl, 2 mM MgCl₂, 5% Glycerol, 2 mM DTT, 0.2 mM ATP, pH 7.6. The fractions containing purified untagged VPS4B were pooled, flash frozen and stored at -80 °C.

For purification of GST-VTA1, the clarified lysate was loaded on glutathione-agarose resin and incubated for 1 hr at 4°C with rotation. The resin was washed with 10 volumes of lysis buffer and eluted in 20 mM HEPES-KOH, 250 mM KCl, 20 mM Glutathione, 5% Glycerol, 2 mM DTT, pH 7.6. Eluate was treated with PreScission protease to cleave the N-terminal GST tag during dialysis into 20 mM HEPES-KOH, 25 mM KCl, 2 mM DTT, pH 7.7 overnight. Protein was loaded onto a MonoQ 5/50GL (Cytiva) pre-equilibrated in dialysis buffer, washed with 3 column volumes of 3% high salt buffer (20 mM HEPES-KOH, 1 M KCl, 2 mM DTT, pH 7.6) and eluted in a gradient 3%-25% of high salt buffer over 25 column volumes. Protein-containing fractions were concentrated using an Amicon Ultra 10K concentrator to ∼0.5 mL, 0.22 μm filtered, and loaded on a Superdex 200 Increase 10/300 GL column (Cytiva) equilibrated with 20 mM HEPES-KOH, 150 mM KCl, 5% Glycerol, 2 mM DTT, pH 7.6. The fractions containing purified untagged VTA1 were pooled, flash frozen, and stored at -80°C.

### In vitro ATPase assays

In vitro ATPase activity of VPS4B mutant enzymes was measured using the NADH-coupled assay on a plate reader (excitation 340 nm, emission 440 nm; Synergy NEO Microplate Reader), similar to described before^37^. Tagless protein constructs were used for VPS4B and VTA1, mixed in a 1:2 molar ratio (200 nM of VPS4B and 400 nM of VTA1). The assay buffer contained 20 mM HEPES-KOH, 25 mM KOAc, 2 mM MgCl_2_, 5 mM (NH_4_)_2_SO_4_, 0.5 mM CaCl_2_, 1 mM TCEP, pH 7.4, supplemented with 0.1 mg/ml BSA and 0.005% vol/vol Triton X-100 for chemical inhibitors testing. Compounds were dissolved in DMSO and diluted in assay buffer and incubated with VPS4B/VTA1 and NADH-coupling reagents for 20 min at room temperature. Reactions were activated by the addition of 1 mM Na_2_ATP. Reactions with no ATP were used as a background control and were subtracted from measured values. Data was analyzed in GraphPad Prism using a modified Michaelis-Menten equation including a Hill coefficient: ATPase rate = (*V_max_x^h^*)/(*K^h^*_1/23_ + (*IC*_50_/*x*)^*h*^) where *V_max_*, is the maximum ATPase rate, ℎ is the Hill coefficient, *x* is the ATP concentration, and *K*_1/2_ is the ATP concentration required to achieve half-maximal ATPase activity.

## Microscopy data quantifications

### Nuclei segmentation

Cell nuclei were segmented from maximum intensity projections of confocal fluorescence microscopy images using the cyto3 model of Cellpose^88^. Cropped nuclei at the image borders, mitotic cells, micronuclei, and dead cells were excluded from the analysis. Detected masks were manually inspected and corrected, if necessary, in Fiji.

### Spatial analysis of centromere distribution

To quantify the spatial positioning of centromeres within defined nuclear ROIs, a custom ImageJ macro was used. Centromere foci were segmented using the ComDet plugin with the following parameters: 2 pxl approximate spot size, 15 SD threshold, segment larger particles. To describe centromere positioning, metrics independent of nuclear size or shape were used. An Exact Distance Transform (EDT) was used to map the distance of every interior pixel to the nearest perimeter edge. For each centromere, the distance to the nuclear border was calculated and then averaged across all centromeres within the nucleus. This value was normalized to the average obtained from the same 200 Monte Carlo simulations to calculate the Border Positioning Index (BPI). Higher BPI values (BPI > 1) indicate more centrally positioned centromeres within the nucleus.

### Colocalization of centromeres with CHMP7, LEMD2, or Emerin

Centromere co-localization with CHMP7, LEMD2, or Emerin was measured on maximum intensity projections of confocal microscopy images, using ComDet plugin (Include larger particles; segment larger particles; approximate particle size: 2.0; intensity threshold (in SD): 10.0 for CREST, CHMP7, LEMD2, Emerin; max distance between colocalized spots: 5.0; join ROIs for intensity of colocalized particles: true) within pre-segmented nuclei.

### Spots detection and colocalization

Colocalization of CHMP7 and γH2AX spots was analyzed using ComDet (Include larger particles; segment larger particles; approximate particle size: 4.0; intensity threshold (in SD): 10.0; max distance between colocalized spots: 5.0; join ROIs for intensity of colocalized particles: true).

### Analysis of nuclear localization of ESCRT-III subunits

For analysis of ESCRT-III subunits nuclear localization upon VSP4 inhibition and depletion was quantified from maximum-intensity projections of confocal microscopy images using Fiji. For VPS4 inhibition nuclei were segmented using the cyto3 model of Cellpose, then camera background intensity was subtracted (100), and fraction of nuclear intensity was measured as ratio of fluorescence intensity within nuclear mask / total intensity. Enrichment of nuclear fraction after VPS4 inhibition was measured as Alk/DMSO-treated ratio. For VSP4 depletion, binary masks of the protein of interest and nuclei (Hoechst channel) were generated in Fiji. The total area occupied by the protein of interest was measured using the Analyze Particles function (size = 0-Infinity; circularity = 0.00-1.00). To quantify nuclear protein localization, the protein mask was intersected with the nuclear mask using the Fiji Image Calculator AND function, and the resulting nuclear protein-positive area was measured. The fraction of protein signal localized to the nucleus was calculated as the nuclear protein-positive area divided by the total protein-positive area.

---

**Figure S1.**
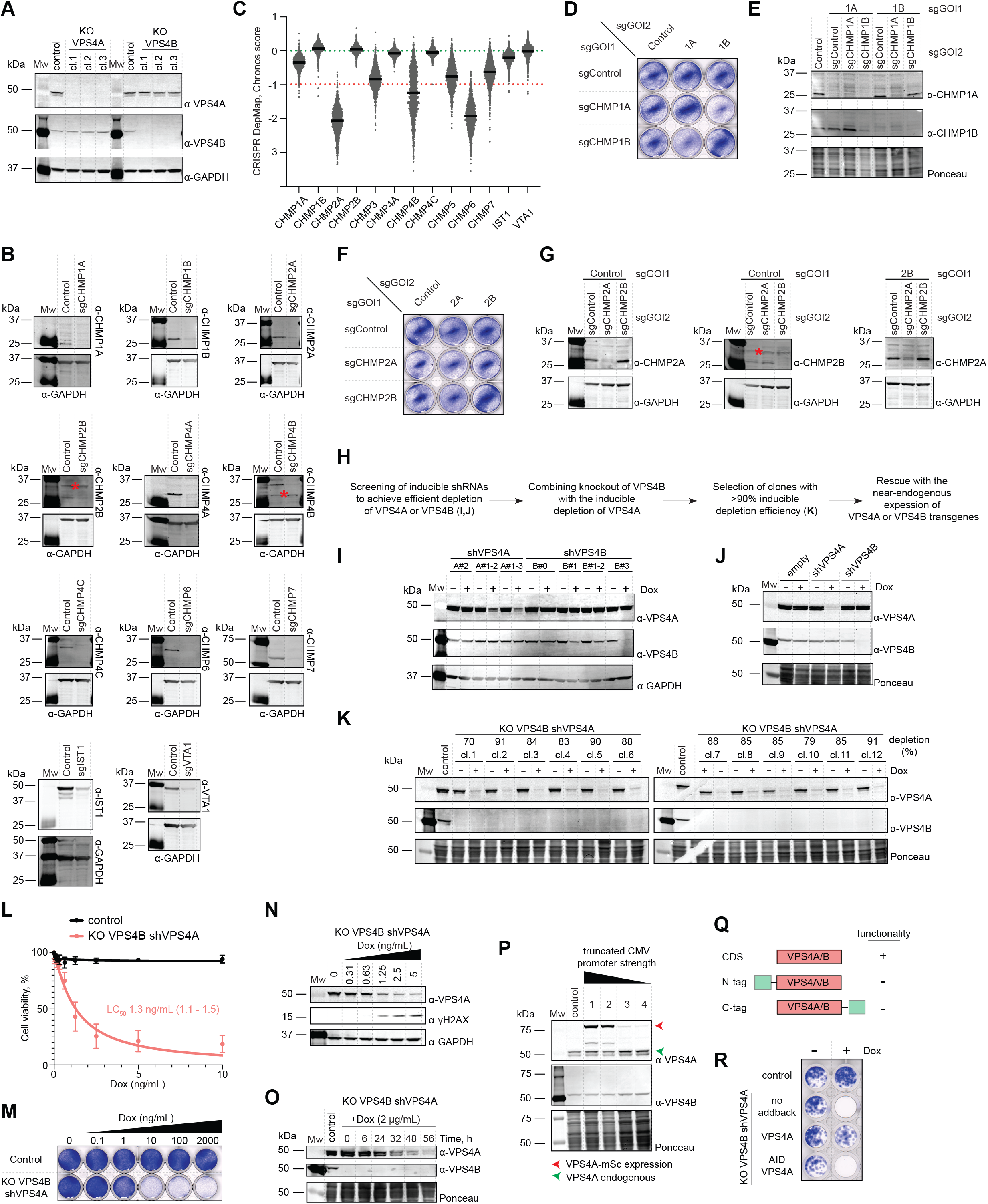
(A) Immunoblot analysis of knockouts of individual VPS4 paralogs. Three clones for each paralog knockout are shown. α-GAPDH was used as a loading control. (B) Immunoblot analysis of CRISPR-mediated depletion of ESCRT-III subunits. Samples were collected 48 h post-selection of transfected cells. A red star indicates a non-specific band (for α-CHMP2B and α-CHMP4B immunoblots). (C) DepMap Chronos scores plotted for ESCRT-III subunits and VTA1. Each gray dot represents a cell line. Dashed lines show median Chronos scores across essential genes (−1, red) and non-essential (0, green). (D) Analysis of cell viability following combined CRISPR-mediated depletion of CHMP1A and CHMP1B paralogs. Abbreviation: GOI, gene-of-interest. (E) Immunoblot analysis of combined CRISPR-mediated depletion of CHMP1A and CHMP1B paralogs. (F) Analysis of cell viability following combined CRISPR-mediated depletion of CHMP2A and CHMP2B paralogs. (G) Immunoblot analysis of combined CRISPR-mediated depletion of CHMP2A and CHMP2B paralogs. A red star indicates a non-specific band (for α-CHMP2B immunoblot). (H) Schematic for the development VPS4 conditional depletion-rescue system (see Methods). (I) Immunoblot analysis for screening of shRNAs targeting VPS4A or VPS4B in HeLa cells. Depletion was induced by addition of Dox (2 µg/mL for 48 h). (J) Immunoblot analysis of optimized shRNAs for depletion of VPS4A or VPS4B. Depletion was induced by addition of Dox (2 µg/mL for 48 h). (K) Immunoblot analysis of shVPS4A in KO VPS4B cells. Screening of clones for efficient depletion. Depletion was induced by addition of Dox (2 µg/mL for 48 h). (L) Analysis of viability of control cells (parental HeLa; black) and KO VPS4B shVPS4A (red) measured 72 h after doxycycline (Dox) addition. Values were normalized to the 0 ng/mL Dox treated conditions (100%). Points represent mean values ± SD (n = 4, 3 technical repeats each). LC_50_ with 95% CI (shown in the parentheses) is calculated based on a sigmoid inhibition curve fit with variable slope. Fit parameters are (Hill coefficients (*h*)=-1.149; R^2^=0.94). (M) Analysis of cell viability of KO VPS4B shVPS4A cells upon Dox-mediated VPS4A depletion (indicated concentrations; 72 h). (N) Immunoblot analysis of KO VPS4B shVPS4A cells shows that decreasing VPS4A expression induces dose-dependent (Dox) accumulation of γH2AX. Depletion of approximately half of the remaining VPS4A (at ∼1 ng/mL of Dox) leads to the cell death; depletion was induced by Dox addition (indicated concentrations; samples were collected at 48 h). (O) Immunoblot analysis of KO VPS4B shVPS4A cells shows the time course of VPS4A depletion. Depletion was induced by Dox addition (2 µg/mL; cells were treated for indicated time). (P) Immunoblot analysis of constitutively expressed tagged VPS4A transgenes, using truncated CMV promoters. A red triangle points to the expressed transgenes; a green triangle points to the endogenous protein. Abbreviation: CDS, coding sequence. (Q) Schematic of VPS4 rescue constructs. Both N- and C- terminally tags on VPS4A and VPS4B compromise their functionality. (R) Analysis of cell viability in cells expressing AID-tagged VPS4A construct using the developed VPS4 conditional depletion system. Cells were grown for 9 days.

**Figure S2.**
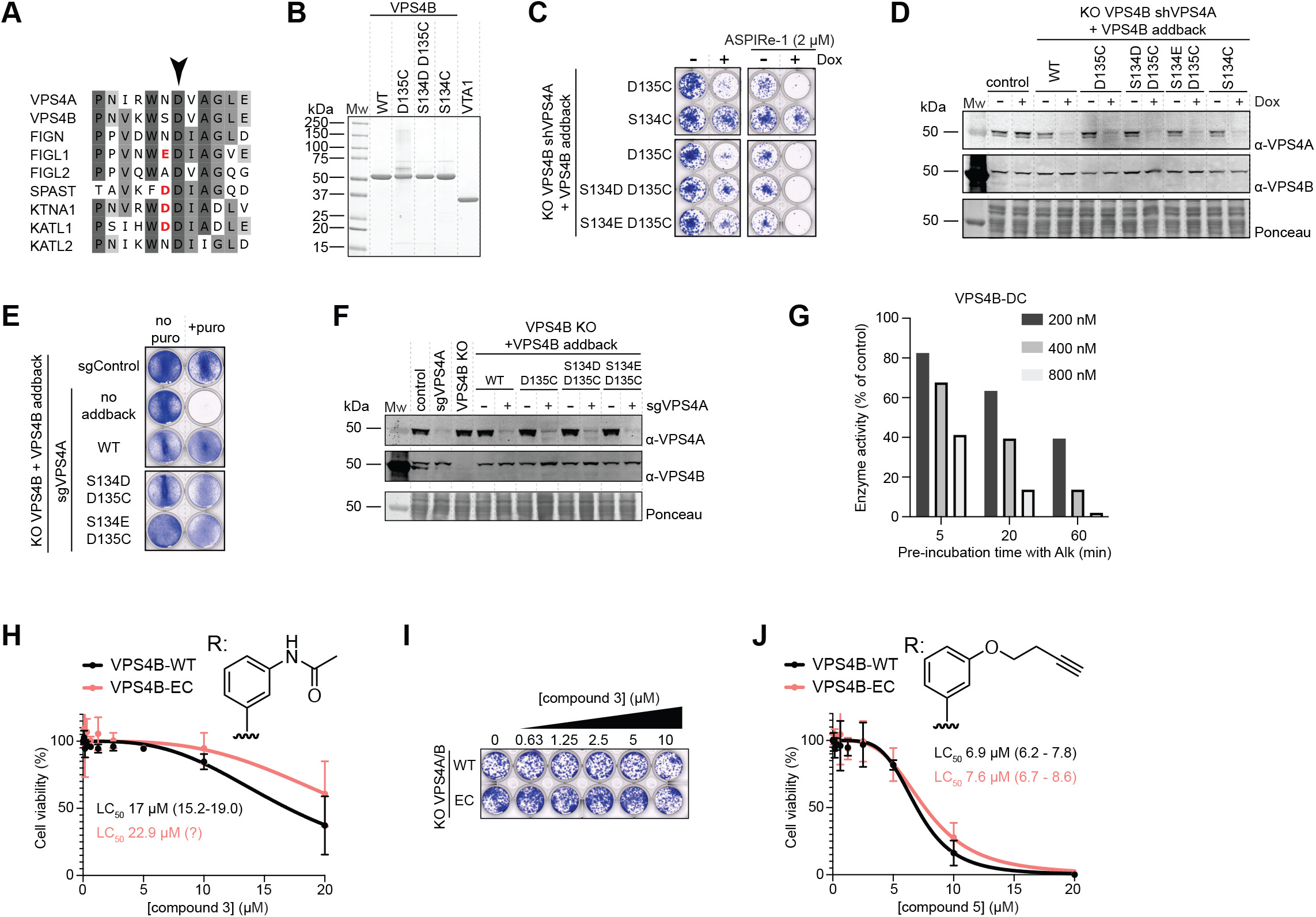
(A) Sequence alignment of the meiotic clade of AAA enzymes, human proteins are shown. The position of the cysteine mutation is a Asp135 in human VPS4B (black arrow). A negatively charged amino acid preceding Asp135 residue is highlighted (red). (B) SDS-PAGE of purified recombinant VPS4B and VTA1 proteins (following cleavage of the purification tag); gel is stained with Coomassie Blue. (C) Analysis of VSP4B cysteine mutants inhibition by ASPIRe-1 in a VPS4 conditional depletion system. Depletion was induced by doxycycline addition (2 µg/mL; cells were grown for 8 days). (D) Immunoblot analysis of KO VPS4B shVPS4A cells stably expressing VPS4B-WT or cysteine mutants. Depletion was induced by doxycycline addition (2 µg/mL for 48 h). Control indicates parental HeLa cells. (E) Analysis of cell viability upon VPS4A/B double knockout using CRISPR/Cas9, rescued with the indicated VPS4B constructs. Cells were fixed and stained (Crystal violet; 48 h post-selection). (F) Immunoblot analysis of VPS4A/B double knockout cells rescued with the indicated VPS4B mutants. Cells were collected 48 h post-selection. Control indicates parental HeLa cells. (G) Analysis of time-dependent inhibition of VPS4B-DC ATPase activity by ASPIRe-Alk is consistent with covalent inhibitor binding. The assay was done at 1 mM of ATP. (H) Analysis of cell viability of VPS4B-WT (black) and VPS4B-EC (red) treated with compound 3 (72 h; 96-well plate assays). Values were normalized to DMSO-treated conditions (100%). Points represent mean values ± SD (n = 3, 3 technical repeats each). LC_50_ with 95% CI are shown in parentheses. Values were estimated based on sigmoid inhibition curve fits with variable slope; the confidence interval for the VPS4B-EC LC_50_ could not be determined. Hill coefficients (*h*) and R^2^: for VSP4B-WT (−3.06; R^2^=0.86), for VSP4B-EC (−3.28; R^2^=0.53). (I) Colony formation assay of VPS4B-WT and VPS4B-EC cells treated with compound 3 (cells were grown for 8 days). (J) Analysis of cell viability of VPS4B-WT (black) and VPS4B-EC (red) treated with compound 5 (72 h; 96-well plate assays). Values were normalized to DMSO-treated conditions (100%). Points represent mean values ± SD (n = 3, 3 technical repeats each). LC_50_ with 95% CI are shown in parentheses. Values were estimated based on sigmoid inhibition curve fits with variable slope. Hill coefficients (*h*) and R^2^: for VSP4B-WT (−4.50; R^2^=0.95), for VSP4B-EC (−3.67; R^2^=0.94).

**Figure S3.**
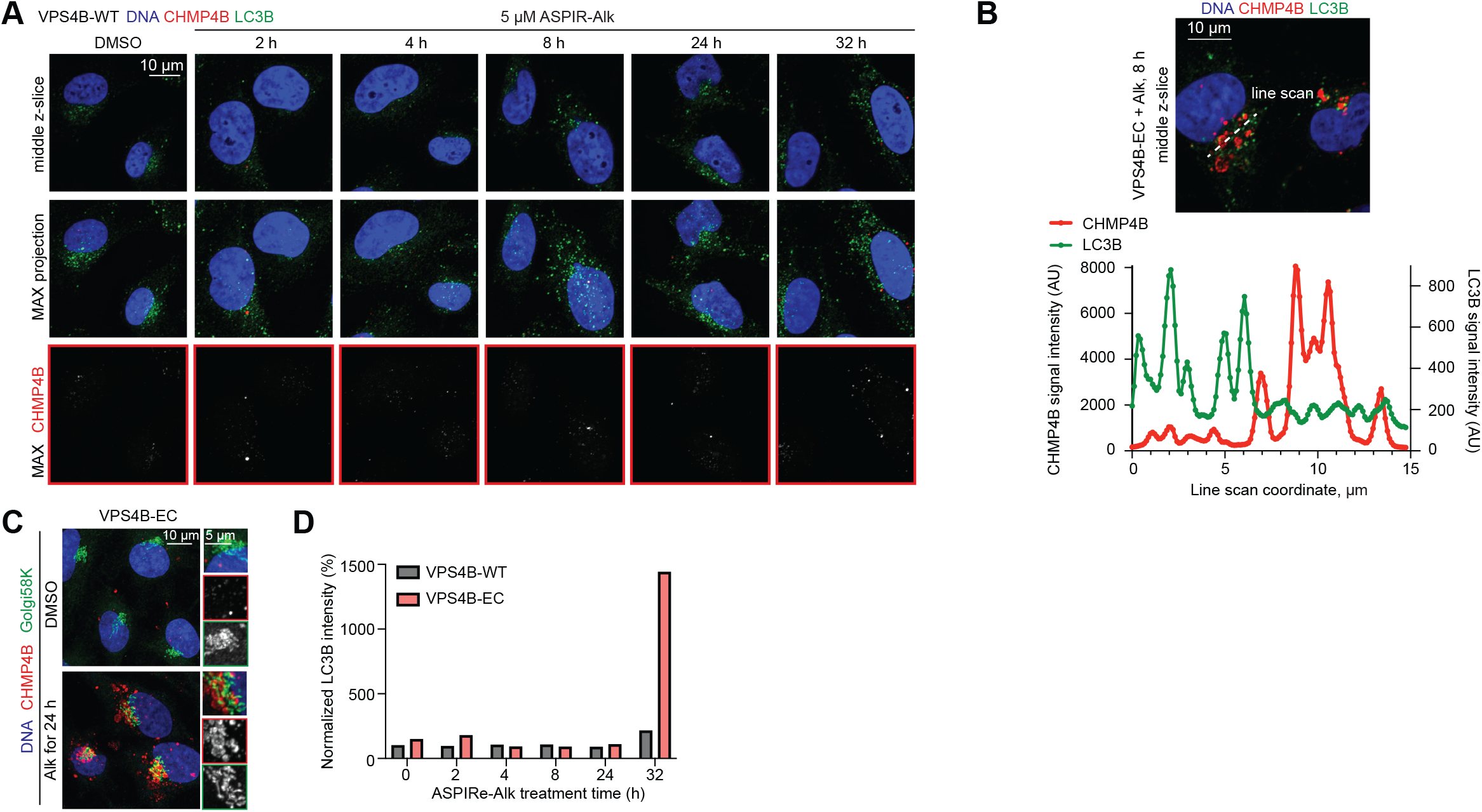
(A) Representative immunofluorescence images of CHMP4B and LC3B following treatment of VPS4B-WT with ASPIRe-Alk (5 µM; indicated time). Middle z-slice and maximum intensity projection are shown. Scale bar, 10 µm. (B) Line scan analysis of CHMP4B and LC3B following VPS4B-EC cells treatment with ASPIRe-Alk (8 h time point; the same image is shown as in Figure 3I). Scale bar, 10 µm. Intensity profiles of CHMP4B and LC3B along the line scan of 5 pxl width (bottom). (C) Immunofluorescence image of CHMP4B and Golgi58K following treatment of VPS4B-EC cells with ASPIRe-Alk (5 µM for 24 h); scale bar, 10 µm. Magnified boxed region is shown on the right side. From top to bottom: merged image, CHMP4B channel, Golgi58K channel. Scale bar, 5 µm. (D) Analysis of LC3B accumulation following VPS4B-EC cells treatment with ASPIRe-Alk. Quantification was performed using part of the data shown in Figure 3L. Average values of two experiments are shown.

**Figure S4.**
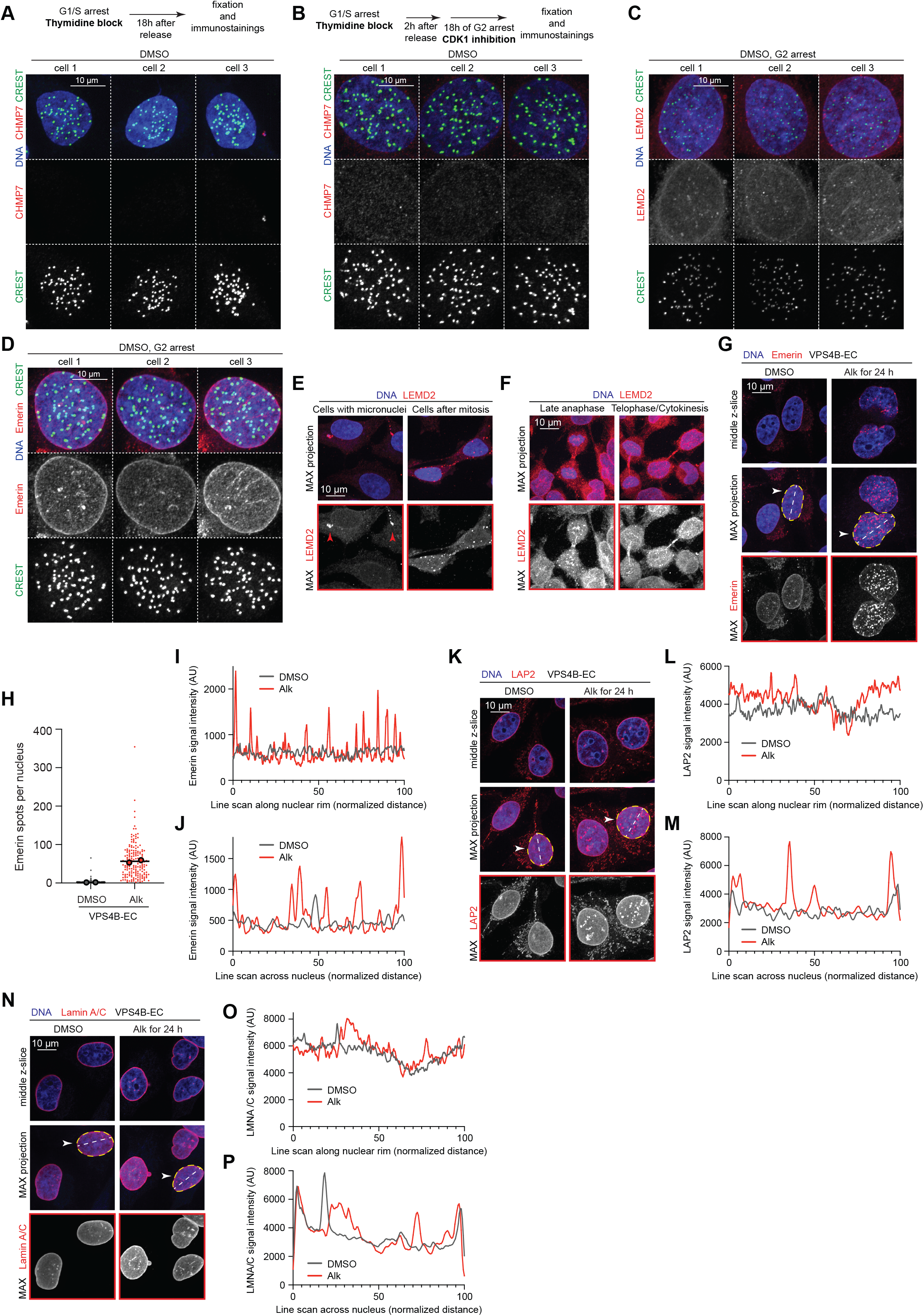
(A) Representative immunofluorescence images of CHMP7 and centromeres in VPS4B-EC cells treated with DMSO. Cells were released from a thymidine block, and progressed through mitosis (A) or were arrested in G2 for 18h (B). Maximum intensity projections of z-stacks are shown; scale bars, 10 µm. (C) and (D) Immunofluorescence images of LEMD2, Emerin and centromeres in VPS4B-EC cells treated with DMSO and arrested in G2 for 18h. Maximum intensity projections of z-stacks are shown; scale bars, 10 µm. (E) Immunofluorescence images of LEMD2 accumulation in micronuclei and cytoplasmic LEMD2 spots in cells after mitosis. Arrowheads point to micronuclei. Scale bar, 10 µm. (F) Immunofluorescence images of LEMD2 accumulation on chromatin at the end of mitosis. Scale bar, 10 µm. (G) Immunofluorescence images of Emerin in VPS4B-EC cells treated with DMSO or ASPIRe-Alk (5 µM for 24 h). Middle z-slices and maximum intensity projections are shown. Cells used for line scan analysis are indicated (arrowhead and dotted line). Scale bar, 10 µm. (H) Analysis of the number of Emerin spots per nucleus following treatment of VPS4B-EC cells with ASPIRe-Alk (5 µM for 24 h). Mean values are shown; small dots represent individual nuclei; big dots are mean values of two datasets (total number of cells DMSO: 301; Alk: 175). (I) and (J) Line scan analysis (5 pxl width) of Emerin signal intensity along the nuclear rim and across the nucleus in DMSO (gray) and Alk-treated cells (red). (K) Immunofluorescence images of LAP2 in VPS4B-EC cells treated with DMSO or ASPIRe-Alk (5 µM for 24 h). Middle z-slices and maximum intensity projections are shown. Cells used for line scan analysis are indicated (arrowhead and dotted line). Scale bar, 10 µm. (L) and (M) Line scan analysis (5 pxl width) of LAP2 signal intensity along the nuclear rim and across the nucleus in DMSO (gray) and Alk-treated cells (red). (N) Immunofluorescence images of Lamin A/C in VPS4B-EC cells treated with DMSO or ASPIRe-Alk (5 µM for 24 h). Middle z-slices and maximum intensity projections are shown. Cells used for line scan analysis are indicated (arrowhead and dotted line). Scale bar, 10 µm. (O) and (P) Line scan analysis (5 pxl width) of Lamin A/C signal intensity along the nuclear rim and across the nucleus in DMSO (gray) and Alk-treated cells (red).

**Figure S5.**
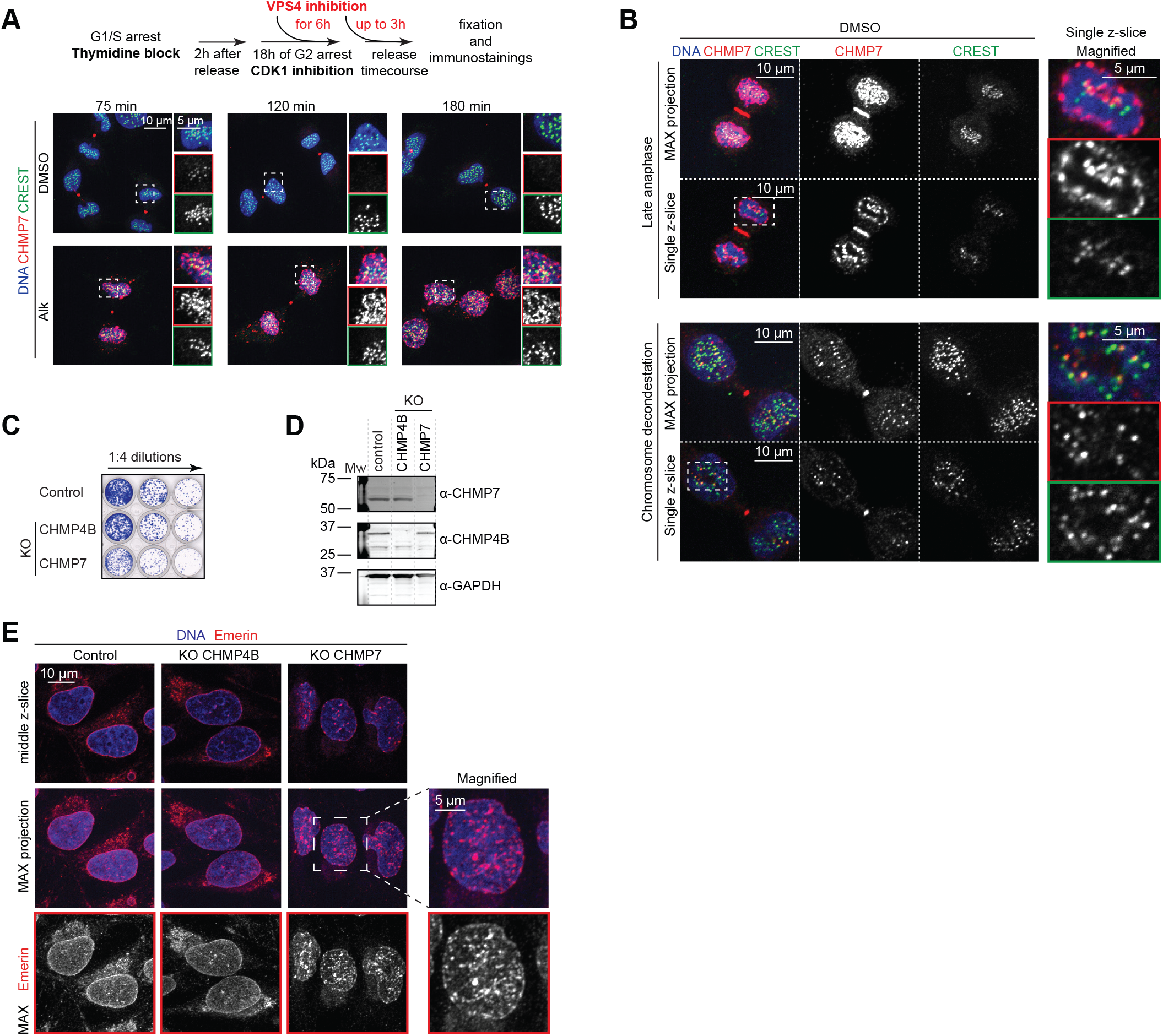
(A) Representative immunofluorescence images of CHMP7 and centromeres (CREST) following treatment of VPS4B-EC cells with ASPIRe-Alk (5 µM for 6 h). Cells were released from a G2 arrest (18 h) and fixed at indicated time points to capture nuclear envelope reformation and chromosome decondensation. Magnified boxed region is also shown. From top to bottom: merged image, CHMP7 channel, CREST channel. Scale bars are 10 µm and 5 µm, as indicated. (B) Immunofluorescence images of CHMP7 and centromeres (CREST) following treatment of VPS4B-EC cells with DMSO. Cells were released from a G2 arrest. Maximum intensity projection and single z-slice are shown; scale bars, 10 µm. Magnified boxed region is also shown; scale bar, 5 µm. From top to bottom: merged image, CHMP7 channel, CREST channel. (C) Analysis of viability of Control (parental HeLa), KO CHMP4B, and KO CHMP7 cells. Colony formation assay (cells were grown for 7 days). (D) Immunoblot analysis of KO CHMP4B and KO CHMP7 cells. Control indicates parental HeLa cells. (E) Immunofluorescence images of Emerin in Control (parental HeLa), KO CHMP4B, and KO CHMP7 cells. Middle z-slices and maximum intensity projections are shown. Scale bar, 10 µm. Magnified boxed region is also shown; scale bar, 5 µm.

**Figure S6.**
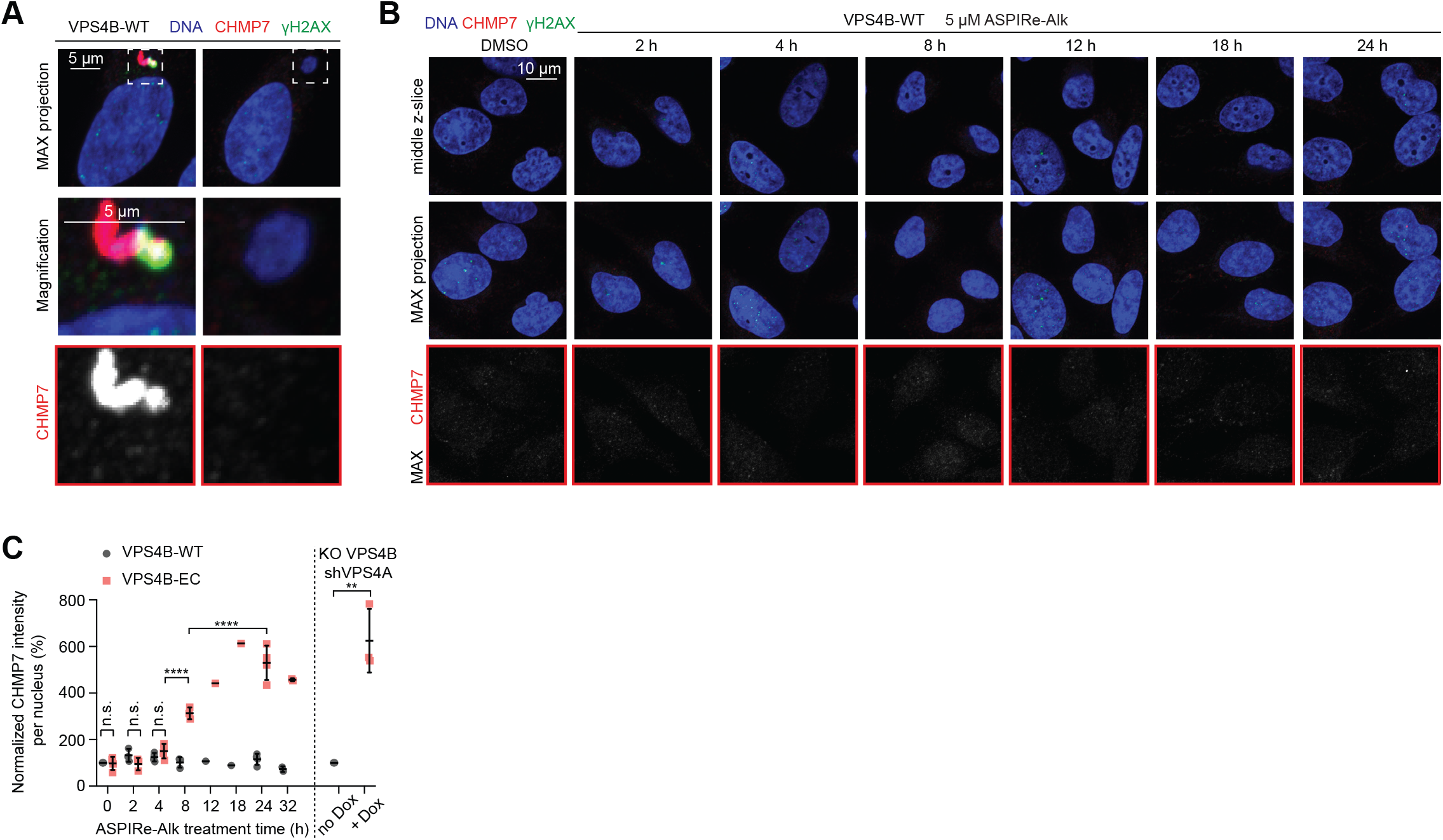
(A) Immunofluorescence images of CHMP7-positive and CHMP7-negative micronuclei in DMSO-treated VPS4B-WT cells. Magnified boxed regions are also shown; scale bars, 5 µm. (B) Immunofluorescence images of CHMP7 and γH2AX following treatment of VPS4B-WT cells with ASPIRe-Alk (5 µM, indicated time). Middle z-slices and maximum intensity projections are shown; scale bar, 10 µm. (C) Time-dependent accumulation of CHMP7 in the nucleus following VPS4B-EC treatment with ASPIRe-Alk (5 µM, time course). The CHMP7 spot intensities per nucleus were quantified. Values were normalized to the DMSO-treated VPS4B-WT condition (100%). Mean values ± SD are shown (points represent independent experiments; >100 cells per point). One way ANOVA with Šídák’s multiple comparisons test. ****p<0.0001; n.s., not significant. For the VPS4 depletion experiment, the same dataset as in Figure 4D was used. Unpaired t test; **p=0.0027.

**Figure S7.**
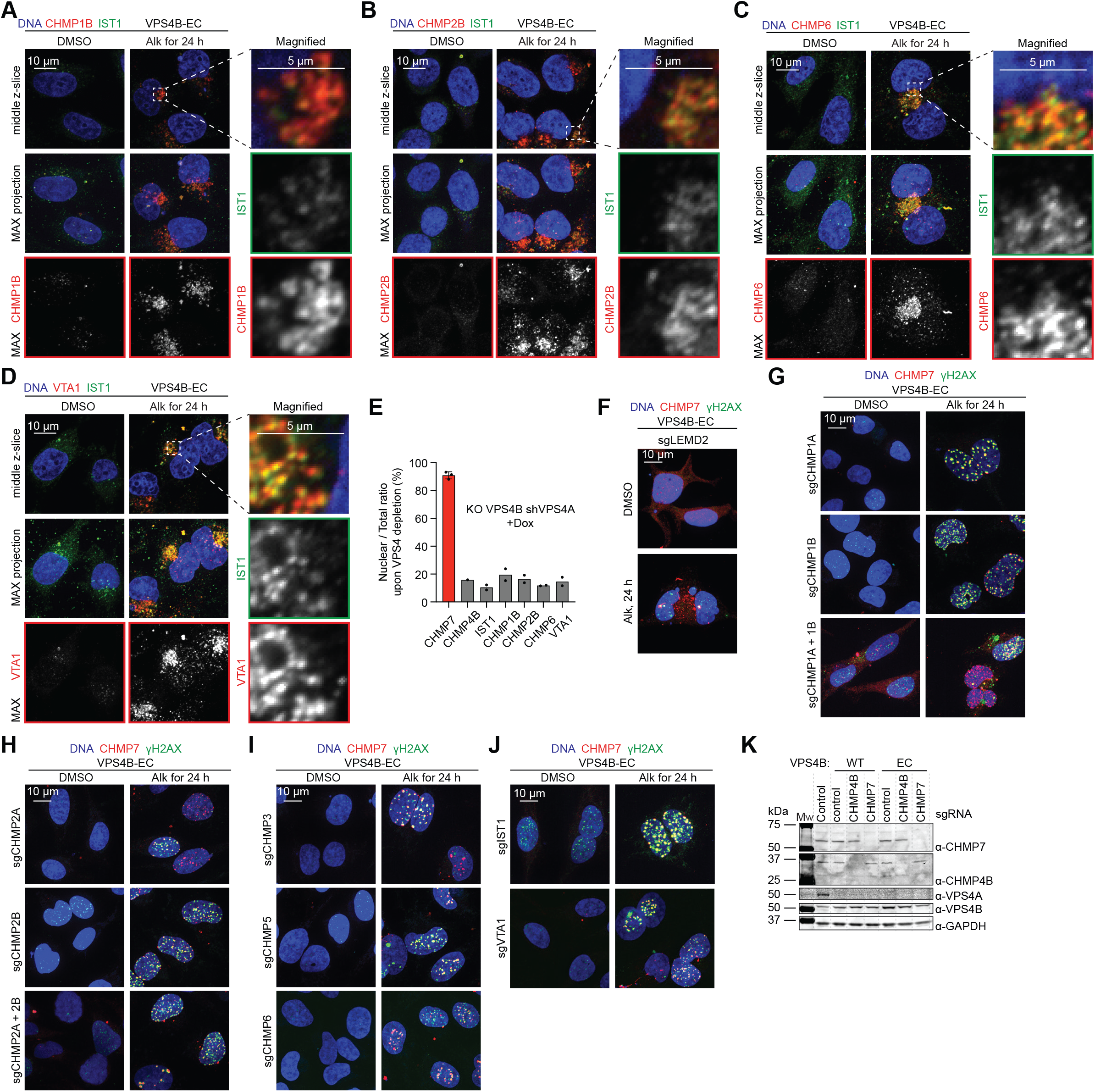
(A-D) Immunofluorescence images of CHMP1B, CHMP2B, CHMP6, VTA1, and IST1 following VPS4B-EC cells treatment with ASPIRe-Alk (5 µM for 24 h). Middle z-slices and maximum intensity projections are shown; scale bar, 10 µm. Magnified boxed regions are also shown; scale bar, 5 µm. (E) Analysis of nuclear accumulation of the indicated ESCRT-III proteins following VPS4 depletion induced with Dox (2 µg/mL for 48 h). Bars are mean values ± SD (points represent independent experiments). (F-J) Immunofluorescence images of CHMP7 and γH2AX following VPS4B-EC cells treatment with ASPIRe-Alk (5 µM for 24 h) combined with depletion of LEMD2 (F), indicated ESCRT-III proteins, or VTA1. Maximum intensity projections of z-stacks are shown; scale bar, 10 µm. (K) Immunoblot analysis of VPS4B-WT and VPS4B-EC cells following CHMP4B or CHMP7 depletion. Control indicates parental HeLa cells.

